# Interpreting and Validating a Deep Learning Model Predictive of Spatial Morphologic-Molecular Patterns in Lung Adenocarcinoma, Using Ground Truth Immunohistochemistry Images

**DOI:** 10.64898/2026.04.20.719723

**Authors:** Vibha R. Rao, Adrienne A. Workman, Scott M. Palisoul, Cassandra J. Limoge, Louis J. Vaickus, George J. Zanazzi, Liang Lu, Xiaoying Liu, Shrey S. Sukhadia

## Abstract

Lung adenocarcinoma (LUAD), the most common subtype of non–small cell lung cancer, exhibits profound histological and molecular heterogeneity. While genomic profiling has identified key oncogenic drivers and immune signatures, its use is limited by cost, technical demands and tissue availability. In addition, spatial transcriptomics provides spatially resolved molecular insights but remains challenging and time-consuming. To address this gap, we developed XpressO-Lung, an explanatory deep learning model that predicts gene expression heterogeneity spatially in tumor and its microenvironment on hematoxylin and eosin based diagnostic (Dx) whole-slide images (WSIs) by learning associations between tissue morphology and the corresponding bulk-transcriptomic data. Utilizing 200 LUAD cases from The Cancer Genome Atlas, XpressO-Lung predicted spatial expression patterns of NAPSA, TP53I3, CD8A, TTF1, KRT7, CDKN2A, FOXO1, KEAP1, RB1 and TP53 on Dx-WSIs with AUCs ranging from 0.64 to 0.92. The predicted spatial gene expression patterns aligned with the known morphologic interactions of the tumor and its microenvironment, capturing biological events directly on Dx-WSIs. These spatio-morpho-molecular associations were further validated using immunohistochemistry on an external set of clinical samples at Dartmouth Health, demonstrating concordance between model-predicted spatial patterns and observed histomorphologic features. By coupling predictive performance with spatial interpretability of gene expression on Dx-WSIs, the XpressO-Lung model bridges histopathology and bulk-transcriptomics, enabling explainable spatio-morpho-genomic analyses to advance biomarker discovery, therapeutic stratification and precision oncology in LUAD.

## Introduction

Lung adenocarcinoma (LUAD) is the most prevalent histological subtype of non-small cell lung cancer (NSCLC), accounting for approximately 40% of all lung cancer diagnoses ^1,2^. Originating from glandular epithelial cells in the distal airways, LUAD is characterized by considerable histological and molecular heterogeneity, often presenting with distinct architectural growth patterns such as lepidic, acinar, papillary, micropapillary, complex glandular, and solid subtypes.^3^. This heterogeneity extends beyond the malignant epithelial cells to include the surrounding tumor microenvironment (TME), which comprises stromal, immune, and vascular elements that significantly influence tumor progression and treatment response. Genomic diversity is also prominent, with frequent oncogenic driver mutations in genes such as *EGFR, KRAS, ALK, STK11,* and *TP53* ^4–6^. Current guidelines, including the WHO and CAP lung cancer biomarker reporting protocols, recommend reflex testing for *ALK* and *ROS1* rearrangements, while the NCCN and IASLC-CAP-AMP guidelines further advocate broad molecular profiling to include *EGFR*, *KRAS*, *BRAF*, *MET*, *RET*, *NTRK*, and PD-L1 as part of the routine diagnostic workup^7^.These alterations inform therapeutic strategies, particularly the use of tyrosine kinase inhibitors and immune checkpoint inhibitors, both of which have substantially improved outcomes for subsets of LUAD patients ^8^. Nevertheless, a substantial subset of LUAD patients show unpredictable responses to therapy and widely varying prognoses, underscoring the need for more comprehensive molecular characterization.

A major obstacle to advancing precision medicine in LUAD is the high degree of inter- and intra-tumoral heterogeneity, which includes variation in both cancer cell states and TME composition. Even among patients with identical histological subtypes or driver mutations, differences in gene expression programs can significantly influence tumor progression, immune interactions, and treatment response ^9^. Addressing this complexity requires an expanded biomarker landscape and scalable tools capable of capturing transcriptomic diversity across LUAD cohorts.

Spatial transcriptomics technologies have emerged as powerful tools for mapping gene expression across tumor sections, enabling in situ profiling of both cancer and non-cancer compartments with high resolution ^10^. However, their technical demands, high costs, and limited clinical availability pose significant barriers to widespread implementation. Moreover, these methods often require specialized protocols and non-standard tissue preservation techniques, limiting their applicability to large-scale or retrospective studies ^11^. As a result, there is growing interest in computational approaches that can infer molecular features from more accessible sources, such as routine histopathology slides.

Hematoxylin and eosin (H&E)-stained diagnostic (Dx) whole slide images (WSIs) offer a rich but underutilized source of phenotypic data. Recent developments in computational pathology have leveraged deep learning (DL) models to extract biologically relevant features from WSIs, demonstrating the ability to classify LUAD subtypes ^12^, predict driver mutations ^13^, infer tumor microenvironment composition ^14^, and even approximate transcriptomic signatures ^15^. These models, often based on convolutional neural networks (CNNs) or attention-based architectures, have shown strong performance across several predictive tasks. However, a persistent challenge in these approaches is their lack of interpretability ^16^.

Most DL models in computational pathology function as “black boxes,” providing slide-level predictions without clear explanations of how these decisions are made or which tissue regions contribute most to the outcome ^17,18^. This lack of interpretability remains a major barrier to its clinical application, as it hinders trust in model predictions and complicates biological validation. In the context of gene expression prediction, where transcriptomic signals are influenced by complex histological interactions between tumor cells and the surrounding TME, the inability to localize predictive regions limits insight into the morphological basis of gene regulation and reduces the utility of these models for downstream biomarker discovery. The importance of interpretability in such high-stakes applications has also been recognized at the policy level, with the White House’s National AI Action Plan designating interpretability as a national research priority and emphasizing its critical role in ensuring safe, trustworthy, and clinically meaningful AI deployment in healthcare ^19^.

To address these challenges, we introduce XpressO-Lung, an explanatory deep learning model that predicts spatial gene expression heterogeneity in tumors and TME from H&E-stained Dx WSIs by learning associations between tissue morphology and bulk transcriptomic profiles using our XpressO pipeline ^20^. XpressO-Lung builds upon our prior work in breast ^20^ and melanoma ^21^ cancers, extending the same modular, transparent framework to LUAD and sufficing it further with an external validation using clinical samples at DH and comparing the model-based predictions (heatmaps) with the standard of care (SOC) Immunohistochemistry (IHC) technique performed on the same set.

The XpressO pipeline employs a weakly supervised attention-based multiple instance learning (MIL) approach, enabling it to associate individual image patches with gene expression signals learned from corresponding tissue-based bulk RNA-seq data ^20^. XpressO-Lung produces gene-specific spatial heatmaps that visualize high- and low-attention regions of interest (ROIs) across the tumor and the surrounding TME within the tissue slides, facilitating direct interpretability of gene expression predictions in focused regions of interest, depicting tumor and TME interactions in LUAD. These spatio-morpho-molecular associations from XpressO-Lung were further validated using IHC for all model-predicted biomarkers on an in-house clinical sample-set, demonstrating the concordance between spatially resolved model attention and the observed histo-morphologic patterns in LUAD. Therefore, this ground-truth validated XpressO-Lung bridges digital pathology with bulk-transcriptomics, using weakly supervised deep learning to infer gene expression patterns on H&E-stained Dx-WSIs. This enables trustworthy analysis of gene expression in settings where direct spatial transcriptomic measurements may be technically challenging, expensive, or clinically inaccessible.

Moreover, XpressO-Lung, while showcasing its predictive performance, also pinpoints the specific tumor and TME regions driving each gene expression prediction. This patch-level interpretability, coupled with a rigorously designed training and evaluation strategy and custom analytical methods for exploring high–low gene expression patterns, enables biologically meaningful insights that drives clinical trust and downstream biological discovery.

## Materials and Methods

### Data Collection and Preprocessing

Dx-WSIs of LUAD from 200 patients were obtained from The Cancer Genome Atlas (TCGA-LUAD) via the Genomic Data Commons (GDC) portal ^22^. These WSIs were processed using the segmentation module in XpressO ^20^ which leverages the CLAM framework ^23^ to identify tumor-enriched regions by assigning attention scores to image patches and selecting the most informative regions for analysis (Figure 1). The WSIs were tiled into non-overlapping 256×256 pixel patches at 20× magnification ^24^. The resulting patches were passed through the feature extraction module in XpressO ^20^, which uses a pretrained Vision Transformer (ViT-L/16) ^25^ called Unified Network for Instance-level Representation Learning (UNI) ^26^, trained with DINOv2 ^27^ to extract high-dimensional feature embeddings optimized for histopathological image analysis (Figure 1). These embeddings enabled the downstream classification of gene expression as described in the following sections.

**Figure 1.**
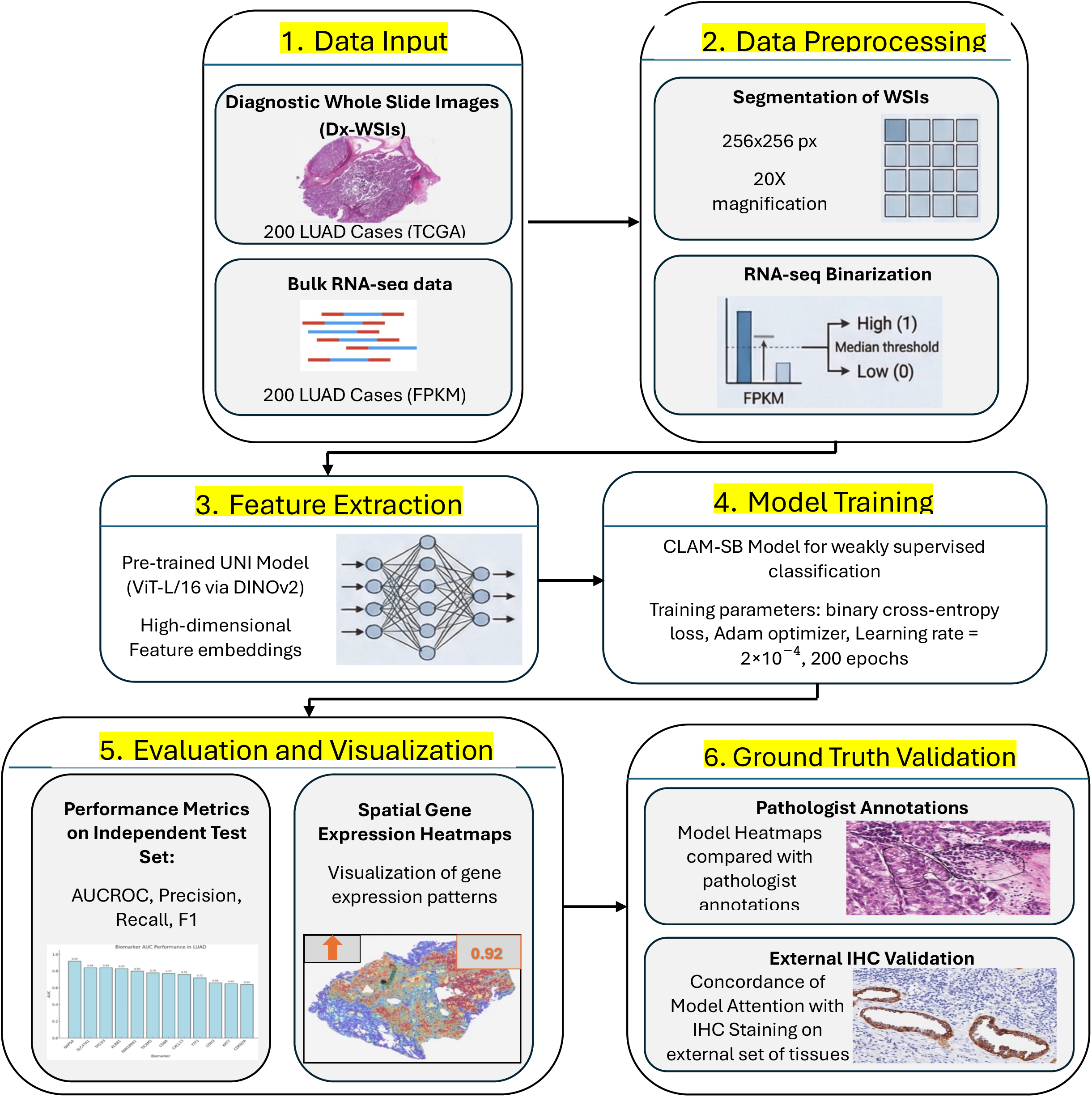
Overview of XpressO Workflow

### Collection and Processing of RNA-seq Data

Corresponding bulk RNA sequencing (RNA-seq) data for the same 200 LUAD patients were downloaded from the TCGA-LUAD portal in the form of fragments per kilobase of transcript per million mapped reads (FPKM). To prepare the dataset for gene expression classification, FPKM values were binarized into “high” and “low” categories using XpressO’s custom binarization script ^20^. For each gene, the median expression value across all patients was used as the threshold with samples with expression values greater than or equal to the median were labelled as high-expression (“1”), while those below the median were labelled as low-expression (“0”). The interquartile range (IQR) of the expression of 10 genes of interest are displayed in Figure 2.

**Figure 2.**
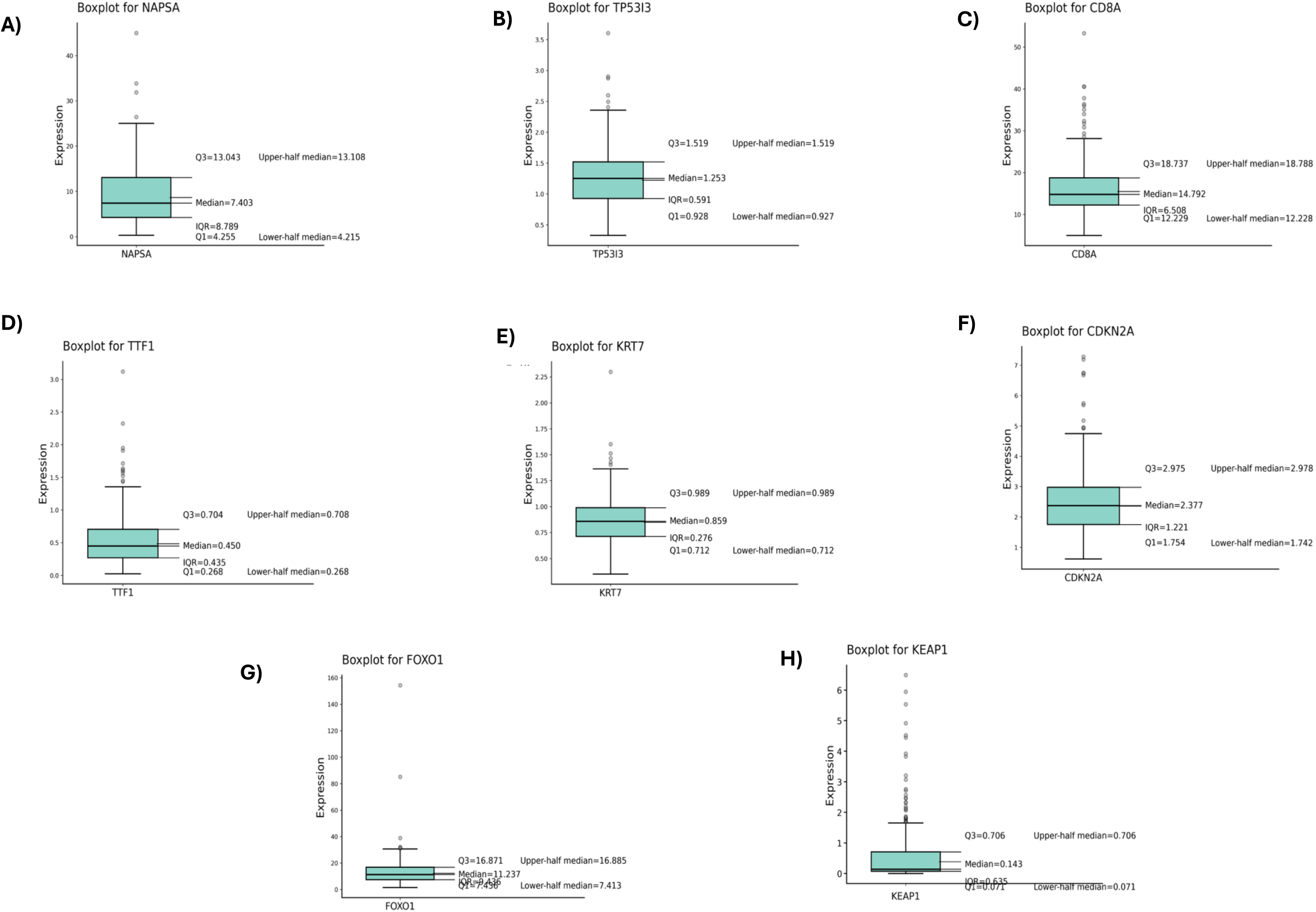
Boxplots summarizing expression levels for ten LUAD markers: (A) NAPSA, (B) TP53I3, (C) CD8A, (D) TTF1, (E) KRT7, (F) CDKN2A, (G) FOXO1, (H) KEAP1, (I) TP53 and (J) RB1 computed across 200 LUAD cases. For each marker, boxplots display the median, first quartile (Q1), third quartile (Q3), interquartile range (IQR), and lower/upper half medians.

Following binarization, the TCGA-LUAD dataset was split into training, internal-validation (used during training to tune hyperparameters and monitor for overfitting) and held-out independent testing subsets using a predefined XpressO script ^20^. Binarization was performed across the full cohort prior to dataset splitting to preserve cohort-wide gene expression variability, enabling the model to learn consistent and robust relationships between histomorphologic features and expression patterns in WSIs. Binary expression labels were assigned at the slide level, allowing each WSI to be categorized as high or low expression for a given gene. These labelled subsets were then used to train and evaluate the deep learning model for gene expression prediction using WSIs.

### Model Training

Gene expression prediction was performed using the classification module in XpressO, which is based on the CLAM-SB architecture ^23^ (Figure 1). For each gene, a weakly supervised model was trained to classify binarized expression status using the top-ranked (k=10) patch embeddings from each tissue-slide in the training set. Training was conducted using binary cross-entropy loss, with the Adam optimizer (learning rate = 2×10^−4^) over 200 epochs. The model was trained to weigh instance-level features using attention-based pooling and output a slide-level prediction for each gene’s expression class.

### Model Evaluation and Metrics

The trained model was evaluated on an held-out independent testing set (i.e. 10% of the total). Evaluation metrics included area under the receiver operating characteristic curve (AUC-ROC), accuracy, precision, recall and F1 scores, along with the 95% confidence intervals for each of them. These metrics were computed separately for each gene (expression) prediction by the model on the independent test set.

### Visualization of the Distribution of High and Low Gene Expression on WSIs

Gene expression heatmaps were generated for the testing set of WSIs using the XpressO’s heatmap module to visualize attention scores derived from the model. For each slide, patch-level attention scores corresponding to class probabilities (p_0 and p_1) were computed, where p_0 represents the predicted class (high or low expression) and p_1 represents the non-predicted class (low or high expression). The top-10 patches with the highest attention scores and the bottom-10 patches with the lowest attention scores were selected to capture regions contributing most and least to the model’s prediction, respectively. The attention scores were subsequently overlaid on the original WSIs and visualized using a color gradient (red to blue), where red denotes high attention, followed by yellow (moderate) and blue (low). These heatmaps were used to spatially localize model-associated regions across tumor and TME areas for each biomarker. Predicted heatmaps were compared with pathologists’ annotations to evaluate alignment between model-derived predictions and established histomorphologic patterns.

### Ground Truth Validation using Immuno-histochemistry (IHC) technique

An external set consisting of 8 tissues (ID 1-8) from Dartmouth Health (DH) were run through XpressO-lung model. For each sample, the gene-specific spatial heatmaps and the probability scores for high or low expression of the relevant genes were generated. In parallel, the IHC staining was performed for the same tissues using the standard protocol at DH. The resultant IHC images were compared with the corresponding spatial heatmaps yielded by XpressO-lung for each gene. These were interpreted manually to establish spatial concordance between IHC-based protein expression and XpressO-lung-based gene expression heatmaps (Figures 3 – 11). This orthogonal validation approach enabled a complementary assessment of model predictions at the tissue, gene and protein level, simultaneously.

**Figure 3.**
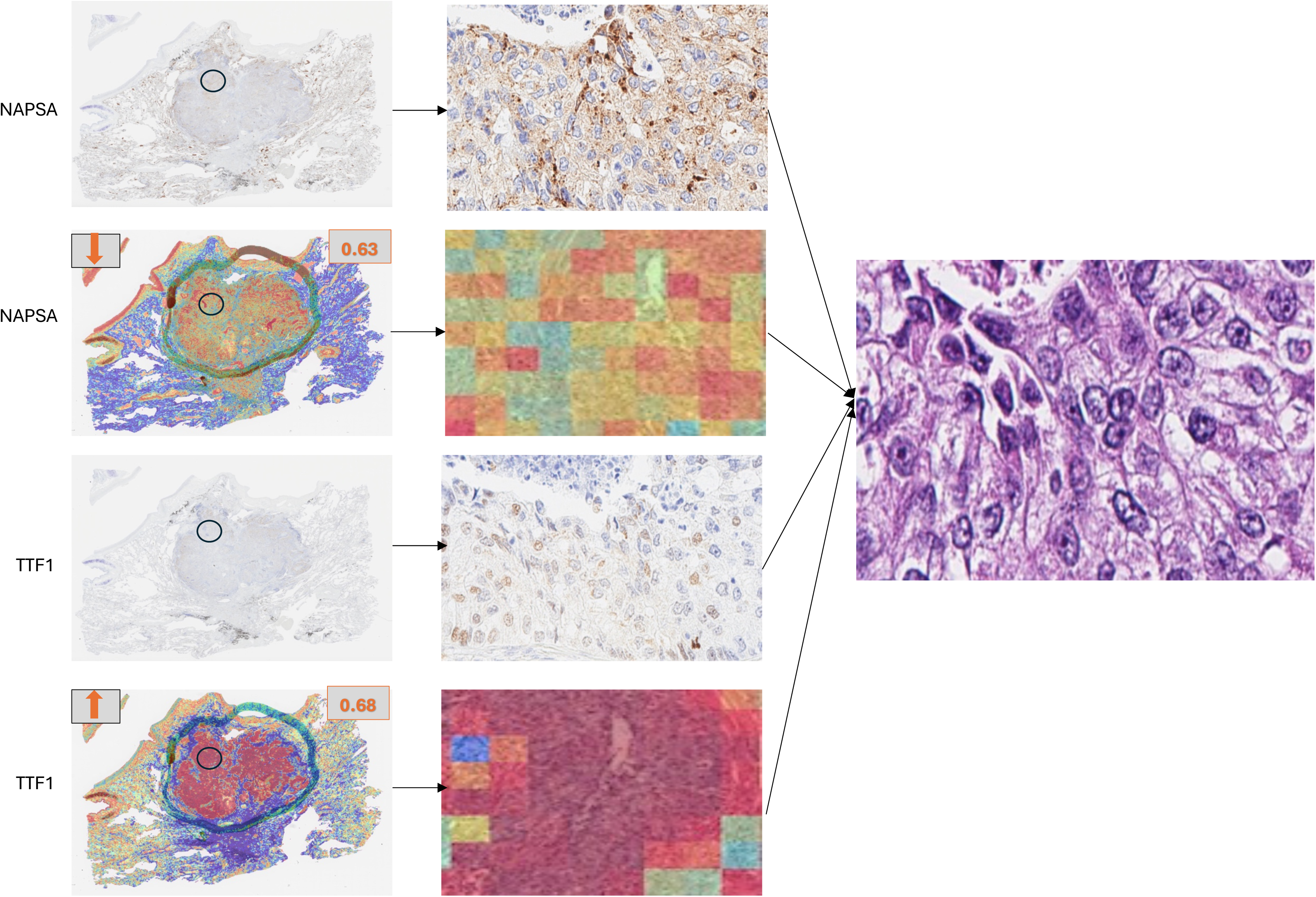
Immunohistochemistry (IHC), model attention maps and corresponding H&E views for NAPSA and TTF1 expression in LUAD tissue sections. Representative cases showing spatial correspondence between IHC staining patterns and underlying histomorphology.

## Results

### Genes of interest

Analysis of bulk RNA-seq profiles from 200 LUAD tumors in the TCGA cohort revealed a broad range of gene expression patterns across patients. Ten genes were selected based on prior literature implicating their varying expression in tumor and TME in LUAD: *CD8A*, *CDKN2A*, *KRT7*, *NAPSA*, *TP53I3*, *TTF1*, *FOXO1*, *RB1*, *KEAP1* and *TP53* (Table 1a). Their fpkm values are depicted in Table 1b. These genes capture key dimensions of LUAD biology, spanning lineage specification, cell cycle control, tumor suppression, and immune context. TTF1 (NKX2-1) ^28–31^, NAPSA (Napsin A) ^32–36^, and KRT7 (CK7) ^37–41^ are canonical IHC markers that define lung adenocarcinoma lineage and are routinely used to distinguish LUAD from squamous and metastatic tumors. CD8A^42–27^ reflects cytotoxic T-cell infiltration and immune activity within the tumor microenvironment. CDKN2A^57–63^, frequently deleted or inactivated in LUAD, drives cell cycle deregulation through loss of p16-mediated control of CDK4/6 activity. Core tumor suppressor pathways are represented by TP53^110^, RB1^109^, and FOXO1^83^, which regulate DNA damage response, cell cycle progression, and apoptotic signaling; their loss or dysfunction is associated with genomic instability and aggressive tumor phenotypes. TP53I3 (PIG3)^64–68^, a p53-inducible gene, links oxidative stress and DNA damage signaling to tumor progression and therapy resistance. KEAP1^108^, a negative regulator of NRF2, is frequently inactivated in LUAD, leading to sustained oxidative stress response, metabolic reprogramming, and resistance to therapy.

**Table 1a:** Genes found to be under- or over-expressed in tumor and/or TME regions in LUAD in the literature.

| Gene Symbol | Full Gene Name | Gene Category | References |
| --- | --- | --- | --- |
| *CD8A* | CD8 Subunit Alpha | TME | 42–47 |
| *CDKN2A* | Cyclin Dependent Kinase<br>Inhibitor 2A | Tumor | 57–63 |
| *KRT7* | Keratin 7 | Tumor | 37–41 |
| *NAPSA* | Napsin A Aspartic Peptidase | Tumor | 32–36 |
| *TP53I3* | Tumor Protein P53 Inducible | Tumor | 64–68 |
|  | Protein 3 |  |  |
| *TTF1* | Thyroid transcription factor 1 | Tumor | 28–31 |
| *FOXO1* | Forkhead Box O1 | Tumor | 83 |
| *RB1* | Retinoblastoma 1 | Tumor | 109 |
| *KEAP1* | Kelch Like ECH Associated<br>Protein 1 | Tumor | 108 |
| *TP53* | Tumor Protein P53 | Tumor | 110 |

**Table 1b:** Genes found to be under under- or over-expressed in tumor and TME regions in the TCGA-LUAD cohort.

| Genes | Gene category | Mean fpkm | Median fpkm | Min. fpkm | Max. fpkm |
| --- | --- | --- | --- | --- | --- |
| *KRT7* | Tumor | 0.881 | 0.859 | 0.351 | 2.298 |
| *TP53I3* | Tumor | 1.273 | 1.253 | 0.332 | 3.604 |
| *CDKN2A* | Tumor | 2.55 | 2.377 | 0.619 | 7.281 |
| *TTF1* | Tumor | 5.623 | 5.045 | 2.282 | 20.554 |
| *NAPSA* | Tumor | 9.412 | 7.403 | 0.3 | 44.973 |
| *CD8A* | TME | 16.658 | 14.792 | 4.996 | 53.298 |
| *FOXO1* | Tumor | 13.58 | 11.24 | 1.545 | 154.306 |
| *RB1* | Tumor | 5.10 | 4.65 | 2.05 | 14.17 |
| *TP53* | Tumor | 1.30 | 0.89 | 0.13 | 12.83 |
| *KEAP1* | Tumor | 0.66 | 0.142 | 0 | 6.49 |

### Model Evaluation of Gene Expression

The performance metrics for the predicted expression of 10 LUAD biomarkers from the testing set are shown in Table 2.

**Table 2:** Performance metrics for the predicted expression of twelve genes in the test set.

| Biomarker | AUC | AUC CI range | Test error | Accuracy | Precision | Precision CI range | Recall | Recall CI range | F1 | F1 CI range | Gen category |
| --- | --- | --- | --- | --- | --- | --- | --- | --- | --- | --- | --- |
| *NAPSA* | 0.92 | (0.84,1.0) | 0.15 | 0.85 | 0.85 | (0.70, 1.00) | 0.85 | (0.7, 1.0) | 0.84 | (0.69, 1.00) | Tumor |
| *TP53I3* | 0.84 | (0.69,1.0) | 0.15 | 0.85 | 0.85 | (0.70, 1.00) | 0.85 | (0.7, 1.0) | 0.85 | (0.69, 1.00) | Tumor |
| *CD8A* | 0.77 | (0.58,0.96) | 0.3 | 0.7 | 0.7 | (0.50, 0.91) | 0.7 | (0.5, 0.9) | 0.69 | (0.48, 0.90) | TME |
| *TTF1* | 0.72 | (0.47,0.96) | 0.3 | 0.7 | 0.84 | (0.77, 0.91) | 0.7 | (0.5, 0.9) | 0.65 | (0.42, 0.89) | Tumor |
| *KRT7* | 0.65 | (0.38,0.92) | 0.35 | 0.65 | 0.68 | (0.47, 0.88) | 0.65 | (0.45, 0.85) | 0.62 | (0.40, 0.85) | Tumor |
| *CDKN2A* | 0.64 | (0.38,0.89) | 0.35 | 0.65 | 0.65 | (0.43, 0.88) | 0.63 | (0.4, 0.85) | 0.63 | (0.41, 0.85) | Tumor |
| *FOXO1* | 0.81 | (0.61,1.0) | 0.25 | 0.75 | 0.65 | (0.40, 0.91) | 0.725 | (0.55, 0.9) | 0.68 | (0.45, 0.90) | Tumor |
| *KEAP1* | 0.77 | (0.58,0.96) | 0.3 | 0.7 | 0.74 | (0.40, 0.83) | 0.7 | (0.4, 0.8) | 0.68 | (0.39, 0.80) | Tumor |
| *TP53* | 0.79 | (0.61,0.96) | 0.25 | 0.75 | 0.80 | (0.64, 0.96) | 0.725 | (0.55, 0.9) | 0.75 | (0.58, 0.92) | Tumor |
| RB1 | 0.77 | (0.58,0.96) | 0.25 | 0.75 | 0.77 | (0.57, 0.96) | 0.75 | (0.55, 0.95) | 0.76 | (0.56, 0.95) | Tum |

### Visualization and Interpretation of Biomarker Expression

To further investigate the model’s spatial attention and interpretability of ROIs, we examined the attention heatmaps produced by XpressO’s HeatMaps module for test WSIs across biomarkers. Depending on the model’s association of the biomarkers’ expression with the tissue-morphological patterns, a biomarker could be predicted as either high or low-expressed in several WSI-patches, where red-patches indicate regions that influence the model’s prediction the most, followed by yellow and green-patches that have moderate and low influence on that prediction (Figures 3-11 and Supplementary Figures S1-S12). The p_0 and p_1 indicate probabilities for the predicted class (high or low-expression) using top-10 patches and non-predicted class (low or high expression) using bottom-10 patches (Table 3). The heatmaps revealed spatial associations between predicted biomarker expression and histomorphologic features in tumor and TME regions (Figures 3-6 and Supplementary Figures S1-S12). The following subsections illustrate representative examples for individual biomarkers and biomarker pairs, linking predicted expression with corresponding histomorphologic patterns.

**Table 3.** Model-predicted biomarker expression status across TCGA lung cancer samples. For each TCGA case, the model predicts whether the biomarker is in high or low expression state; where pred_0 = Predicted expression class (high/low expression) using the top-10 attended patches; p_0 = Probability of the predicted class using top-10 attention; pred_1 = Predicted expression class (high/low) using the bottom-10 attended patches; p_1 = Probability of the predicted class using bottom-10 attention. Associated figure panels (S1-S12) point to corresponding heatmaps and H&E visualization.

| ID | Biomarker | pred_0 | p_0 | Pred_1 | p_1 | Figures |
| --- | --- | --- | --- | --- | --- | --- |
| 1 | *NAPSA* | low_expression | 0.61 | high_expression | 0.39 | S1-a-i |
|  | *TTF1* | high_expression | 0.73 | low_expression | 0.27 | S1-a-ii |
| TCGA-75-7027 | *NAPSA* | low_expression | 0.62 | high_expression | 0.38 | S1-b-i |
|  | *TTF1* | low_expression | 0.54 | high_expression | 0.46 | S1-b-ii |
| TCGA-55-1592 | *NAPSA* | low_expression | 0.8 | high_expression | 0.2 | S2-a-i |
|  | *TTF1* | high_expression | 0.73 | low_expression | 0.27 | S2-a-ii |
| TCGA-44-5643 | *CD8A* | high_expression | 0.88 | low_expression | 0.17 | S3-a-i |
|  | *KRT7* | high_expression | 0.74 | low_expression | 0.26 | S3-a-ii |
| TCGA-49-AAR3 | *CD8A* | high_expression | 0.61 | low_expression | 0.39 | S4-a-i |
|  | *KRT7* | low_expression | 0.69 | high_expression | 0.31 | S4-a-ii |
| TCGA-69-7973 | *CD8A* | high_expression | 0.82 | low_expression | 0.18 | S4-b-i |
|  | *KRT7* | high_expression | 0.78 | low_expression | 0.22 | S4-b-ii |
| TCGA-44-8119 | *KRT7* | high_expression | 0.53 | low_expression | 0.47 | S5-a-i |
|  | *CDKN2A* | low_expression | 0.76 | high_expression | 0.24 | S5-a-ii |
| TCGA-97-A4M5 | *KRT7* | high_expression | 0.77 | low_expression | 0.23 | S5-b-i |
|  | *CDKN2A* | low_expression | 0.66 | high_expression | 0.34 | S5-b-ii |
| TCGA-55-7907 | *NAPSA* | high_expression | 0.54 | low_expression | 0.46 | S6-b-i |
|  | *CDKN2A* | low_expression | 0.55 | high_expression | 0.45 | S6-b-ii |
| TCGA-55-8506 | *NAPSA* | low_expression | 0.79 | high_expression | 0.21 | S6-a-i |
|  | *CDKN2A* | low_expression | 0.63 | high_expression | 0.37 | S6-a-ii |
| TCGA-97-A4M1 | *NAPSA* | high_expression | 0.91 | low_expression | 0.09 | S7-a-i |
|  | *CDKN2A* | low_expression | 0.75 | high_expression | 0.25 | S7-a-ii |
| TCGA-97-A4M5 | *NAPSA* | high_expression | 0.52 | low_expression | 0.48 | S7-b-i |
|  | *CDKN2A* | low_expression | 0.66 | high_expression | 0.34 | S7-b-ii |
| TCGA-05-4397 | *TP53I3* | high_expression | 0.87 | low_expression | 0.13 | S8-a |
| TCGA-78-7153 |  | low_expression | 0.91 | high_expression | 0.09 | S8-b |
| TCGA-MP-A4SY | FOXO1 | low_expression | 0.98 | high_expression | 0.02 | S9-a |
| TCGA-97-7554 |  | low_expression | 0.85 | high_expression | 0.15 | S9-b |
| TCGA-05-4430 | KEAP1 | low_expression | 0.96 | high_expression | 0.04 | S10-a |
| TCGA-55-7913 |  | low_expression | 0.98 | high_expression | 0.02 | S10-b |
| TCGA-78-7540 | RB1 | low_expression | 0.91 | high_expression | 0.09 | S11-a |
| TCGA-97-7547 |  | low_expression | 0.92 | high_expression | 0.08 | S11-b |
| TCGA-49-AAR2 | TP53 | low_expression | 0.99 | high_expression | 0.01 | S12-a |
| TCGA-62-A46S |  | low_expression | 0.97 | high_expression | 0.03 | S12-b |

#### a NAPSA and TTF1

*NAPSA* (Napsin A) and *TTF1*, two well-established markers of LUAD, were frequently predicted to have expression heterogeneity and exhibited high attention weights within similar tumor regions (Table 3, Supplementary Figures S1, S2). In WSIs TCGA-55-A493 and TCGA-55-1592, *TTF1* was highly expressed while *NAPSA* showed low expression. In WSI TCGA-75-7027, both markers exhibited low expression. Heatmaps highlighted a focused spatial pattern for *TTF1*, consistent with its stable nuclear localization in the bulk adenocarcinoma clone (Figure 3-a,b, Supplementary Figure S1). In contrast, *NAPSA*’s scattered expression appeared more diffuse within tumor regions, reflecting its cytoplasmic granular staining and possible dilution by adjacent normal pneumocytes (Supplementary Figures S1, S2) ^84,85^. Morphologically, high-*TTF1* regions showed cohesive gland-forming tumor nests with uniform nuclear features, while low-*NAPSA* areas often corresponded to poorly differentiated foci lacking clear cytoplasmic granularity (Supplementary Figures S1, S2). These observations are consistent with known histopathologic features and demonstrate the model’s capacity to distinguish subcellular expression patterns directly from WSIs ^86^.

These spatial patterns were further supported by IHC validation. TTF1 immunoreactivity was confined to the nuclei of tumor cells, with positive nuclei concentrated within cohesive gland-forming tumor regions. The heatmaps demonstrated predominantly red regions in these areas, corresponding to a higher probability of high expression (68%), consistent with the observed IHC staining pattern. In contrast, NAPSA immunoreactivity in the cytoplasm of tumor cells, appearing as a coarse granular signal with a patchy and heterogeneous distribution across tumor regions (Figure 3). The corresponding heatmaps showed mixed distribution of red, yellow, and cyan pixels, indicating heterogeneous expression with low expression (63%), partially reflecting the variable IHC staining (Table 4, Figure 3). Overall, TTF1 shows strong spatial concordance between IHC and model predictions, while NAPSA demonstrates more heterogeneous alignment, reflecting variability in tumor cell differentiation and local microenvironment.

**Table 4:** Model-predicted biomarker expression status across external set from DH. For each case, the model predicts whether the biomarker is in high or low expression state; where pred_0 = Predicted expression class (low/high expression) using the top-10 attended patches; p_0 = Probability of the predicted class using top-10 attention; pred_1 = Predicted expression class (high/low) using the bottom-10 attended patches; p_1 = Probability of the predicted class using bottom-10 attention. Associated figure panels (Figure 3-11) point to corresponding heatmaps, IHC and H&E visualization.

| ID | Biomarker | pred_0 | p_0 | Pred_1 | p_1 | Figures |
| --- | --- | --- | --- | --- | --- | --- |
| 1 | *NAPSA* | low_expression | 0.63 | high_expression | 0.37 | 3-a-ii |
|  | *TTF1* | high_expression | 0.68 | low_expression | 0.32 | 3-b-ii |
| 2 | *CD8A* | high_expression | 0.72 | low_expression | 0.28 | 4-a-ii |
|  | *KRT7* | high_expression | 0.7 | low_expression | 0.3 | 4-b-ii |
| 3 | *KRT7* | high_expression | 0.66 | low_expression | 0.34 | 5-a-ii |
|  | *CDKN2A* | high_expression | 0.53 | low_expression | 0.47 | 5-b-ii |
| 4 | *NAPSA* | high_expression | 0.6 | low_expression | 0.4 | 6-a-ii |
|  | *CDKN2A* | high_expression | 0.53 | low_expression | 0.47 | 6-b-ii |
| 5 | *TP53I3* | low_expression | 0.86 | high_expression | 0.14 | 7-a-ii |
| 2 | *FOXO1* | high_expression | 0.54 | low_expression | 0.46 | 8-a-ii |
| 6 | *KEAP1* | high_expression | 0.79 | high_expression | 0.21 | 9-a-ii |
| 7 | *RB1* | low_expression | 0.58 | high_expression | 0.42 | 10-a-ii |
| 8 | *TP53* | low_expression | 0.71 | high_expression | 0.29 | 11-a-ii |

#### b CD8A and KRT7

The predicted expression of *CD8A* was consistently high across the three WSIs evaluated (TCGA-44-5643, TCGA-49-AAR3, and TCGA-69-7973) in the test set (Table 3, Supplementary Figure S3, S4). *KRT7* expression was also predicted to be high in two of these slides (TCGA-44-5643 and TCGA-69-7973) (Table 3, Supplementary Figure S3, S4), with only one case (TCGA-49-AAR3) (Table 3 and Supplementary Figure S3-a) showing low predicted *KRT7*. Notably, in TCGA-44-5643 and TCGA-69-7973, where both *CD8A* and *KRT7* expression were high, the model-generated heatmaps showed spatial co-localization of *CD8A*-positive regions with *KRT7*-positive epithelial nests. Morphologically, this translated to lymphocytic cuffs and infiltrates directly abutting or penetrating keratin-rich tumor islands, a configuration characteristic of an immune-inflamed TME. This observation is consistent with established histopathologic patterns in LUAD, where a *CD8A* rim penetrating *KRT7*-positive nests is indicative of an immune-inflamed TME ^87^.

CD8A immunoreactivity was present in small lymphocytes within the stroma, concentrated along the tumor boundary and adjacent to glandular structures. In contrast, KRT7 was present in normal lung epithelium and in neoplastic cells. CD8A-positive lymphocytes were positioned around KRT7-positive neoplastic cells, consistent with an immune-inflamed tumor microenvironment and matching the co-localized attention patterns observed in the model (Figure 4). The model heatmaps for the same WSI further supported this spatial relationship.

**Figure 4.**
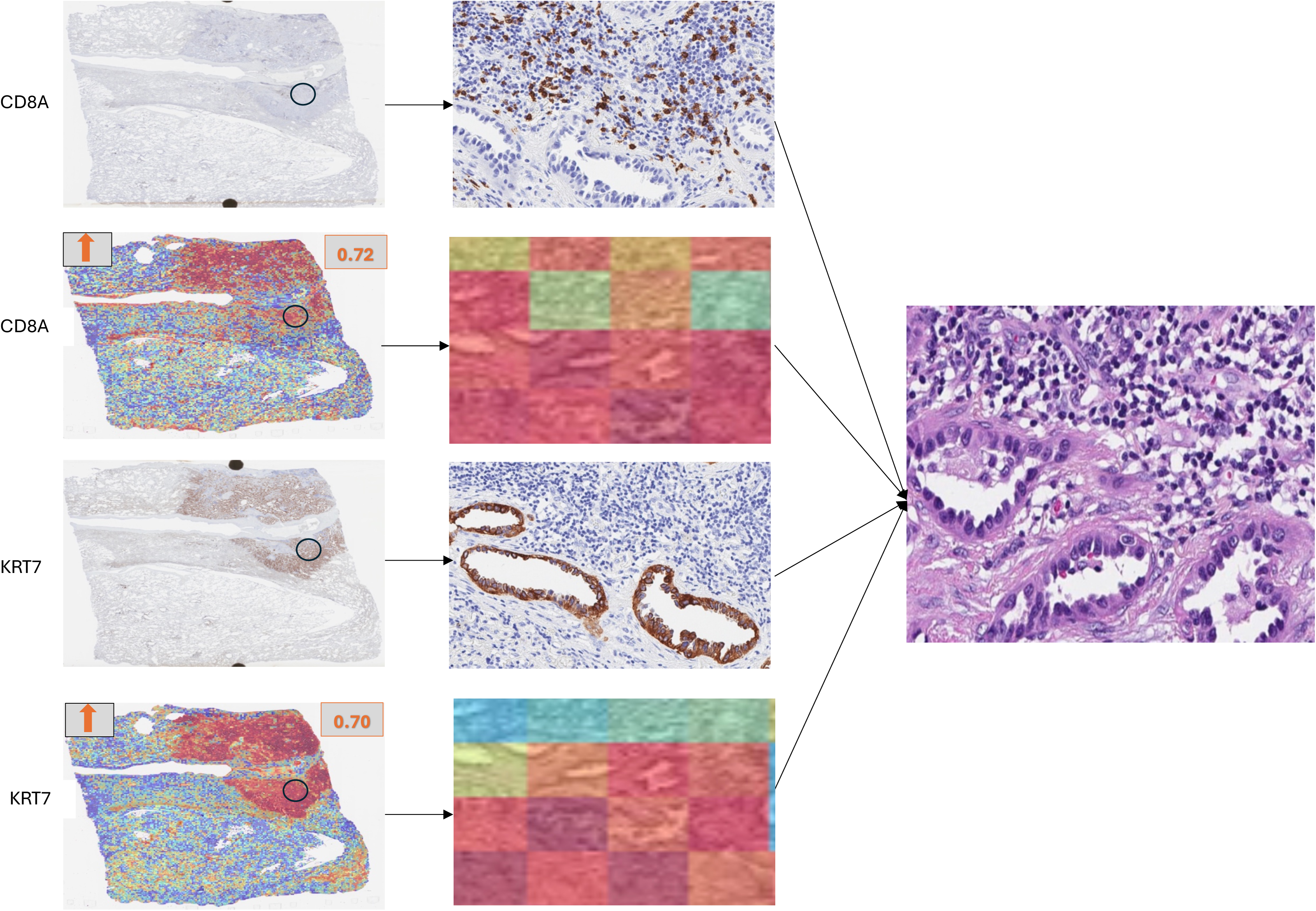
Immunohistochemistry (IHC), model attention maps and corresponding H&E views for CD8A and KRT7 expression in LUAD tissue sections. Representative cases showing spatial correspondence between IHC staining patterns and underlying histomorphology.

Regions enriched with CD8A-positive lymphocytes showed mixed to high attention corresponding to higher expression probability (72%). In contrast, these same regions appeared as blue to cyan in the KRT7 heatmaps, indicating relatively lower expression of KRT7 in the lymphocytes. Conversely, areas corresponding to gland-forming tumor epithelium demonstrated strong red signal in the KRT7 heatmaps (70%) (Table 4 and Figure 4). Together, this highlights spatial complementarity between immune and tumor-associated markers.

#### c KRT7 and CDKN2A

In both test cases analyzed (TCGA-44-8119, TCGA-97-A4M5), *KRT7* expression was predicted to be high, while *CDKN2A* expression was predicted to be low (Table 3 and Supplementary Figure S5). This aligns with previously described histologic patterns in lung adenocarcinoma, where regions that are KRT7-bright but CDKN2A-silent often reflect proliferative LUAD clones with 9p21 loss, a genotype associated with poorer prognosis and potential responsiveness to CDK4/6 or PRMT5-targeted therapies over PD-L1 monotherapy ^88,89^. Morphologically, such regions displayed densely cellular epithelial nests with preserved keratin cytoskeleton but minimal intervening stroma and frequent mitotic figures, supporting a high proliferative index (Supplementary Figure S5).

These patterns were further evaluated by immunohistochemistry. KRT7 demonstrated strong staining outlining glandular tumor structures, with prominent localization along luminal borders. CDKN2A (p16) showed predominantly diffuse brown staining within tumor cells in the evaluated regions, consistent with preserved protein expression (Figure 5). The XpressO-Lung heatmaps for both markers showed heterogeneous distributions with intermixed red, yellow, and blue regions, indicating variable attention across the tissue. KRT7 and CDKN2A showed predicted probabilities of high expression of 66% and 53%, respectively.(Table 4, Figure 5).

**Figure 5.**
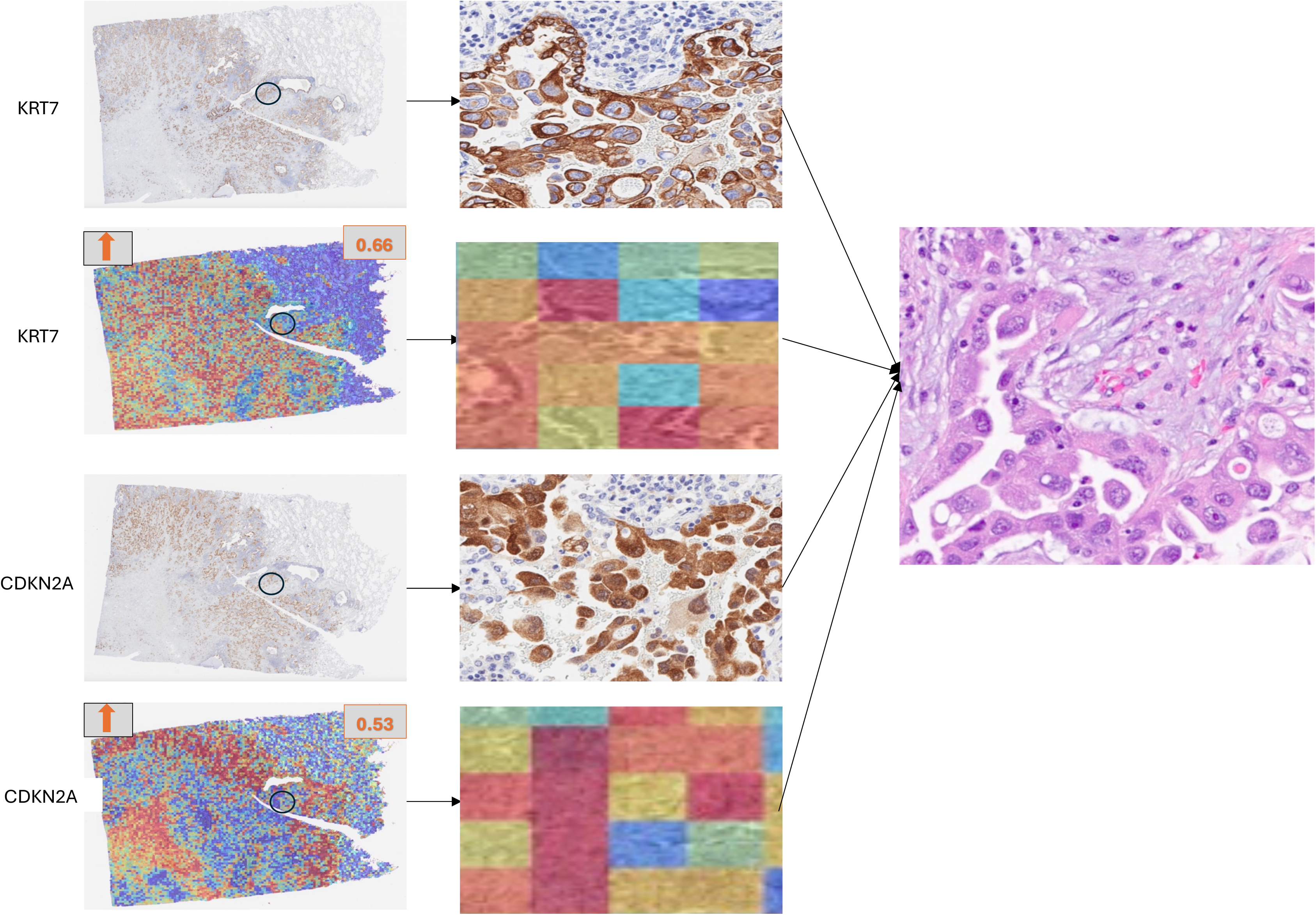
Immunohistochemistry (IHC), model attention maps and corresponding H&E views for KRT7 and CDKN2A expression in LUAD tissue sections. Representative cases showing spatial correspondence between IHC staining patterns and underlying histomorphology.

Overall, these findings reflect heterogeneous but partially concordant spatial patterns between IHC staining and model predictions.

#### d NAPSA and CDKN2A

Across four test tissues (TCGA-55-7907, TCGA-55-8506, TCGA-97-A4M1, TCGA-97-A4M5), *NAPSA* was predicted as high in three and low in one, while *CDKN2A* was consistently low in each tissue, resulting in the most probable pattern of high *NAPSA* coinciding with low *CDKN2A* expression (Table 3, Supplementary Figure S6, S7). This pattern, *NAPSA*–bright yet *CDKN2A*–silent characterizes an alveolar-differentiated but cell-cycle-unleashed LUAD clone, frequently associated with 9p21/MTAP deletions^90^. Morphologically, these regions had abundant, finely vacuolated cytoplasm and centrally placed nuclei, mimicking type II pneumocytes, but arranged in crowded acinar or papillary formations without significant maturation (Supplementary Figure S6, S7).

These patterns were further evaluated by immunohistochemical validation. NAPSA showed strong, dense granular staining in the cytoplasm of tumor cells, consistent with robust expression and alveolar differentiation. In contrast, CDKN2A demonstrated weak-to-moderate, heterogeneous staining within tumor cells, with variable intensity across regions rather than complete absence (Figure 6). The XpressO-Lung heatmaps showed that NAPSA was predominantly represented by cyan and yellow regions with only scattered red areas, indicating moderate but not strongly localized high-expression signal (60%). In contrast, CDKN2A heatmaps showed blue regions corresponding to areas of strong IHC staining within tumor cells, while surrounding stromal regions with lymphocytic infiltration appeared red, indicating higher predicted expression (53%) in areas not corresponding to tumor cell staining (Table 4, Figure 6).

**Figure 6.**
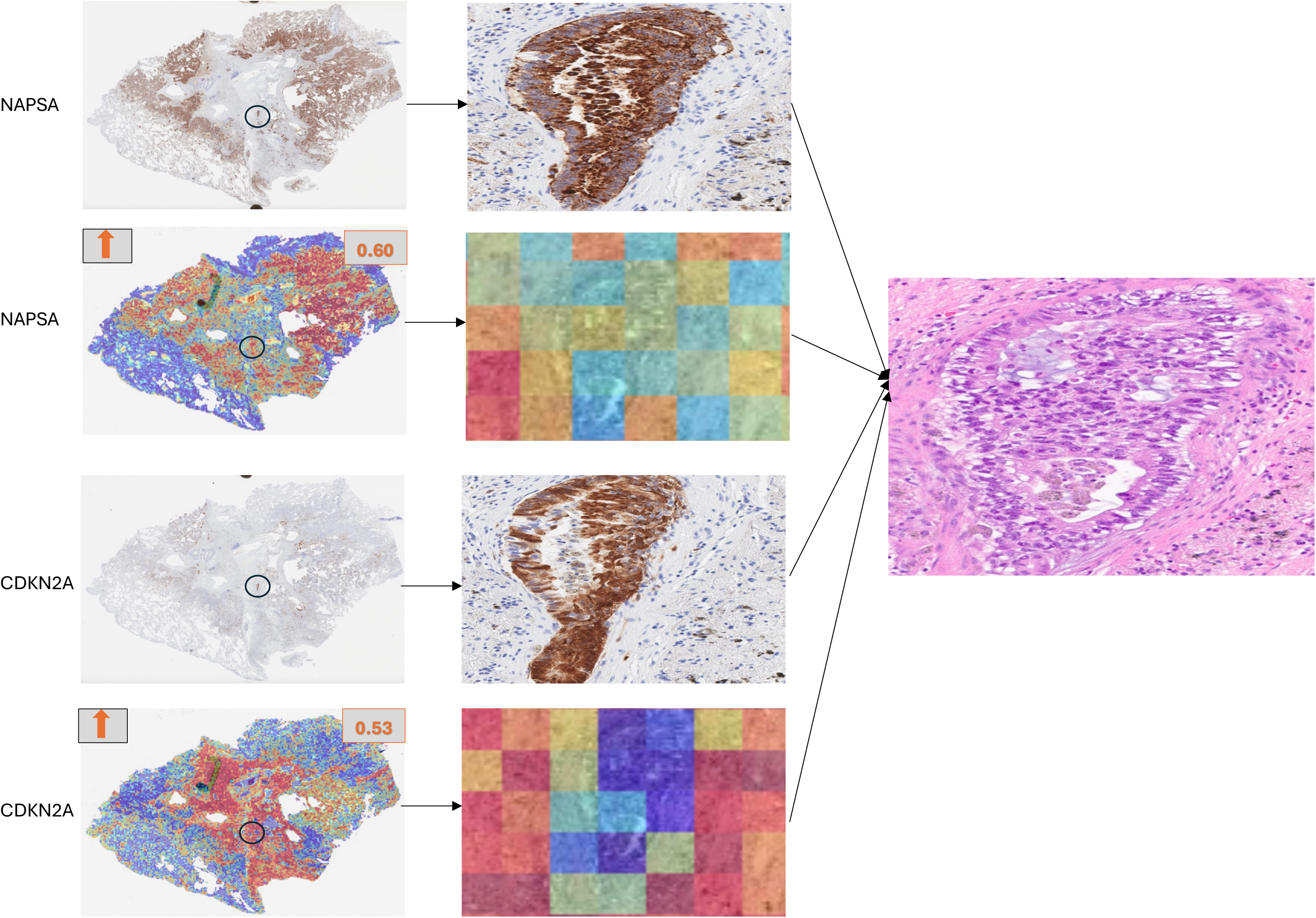
Immunohistochemistry (IHC), model attention maps and corresponding H&E views for NAPSA and CDKN2A expression in LUAD tissue sections. Representative cases showing spatial correspondence between IHC staining patterns and underlying histomorphology.

Overall, this reflects partial alignment for NAPSA and spatial discordance for CDKN2A between IHC staining and model predictions.

#### e TP53I3

*TP53I3*, a transcriptional target of the tumor suppressor TP53, plays a role in oxidative stress response and DNA damage-induced apoptosis ^93^. Predicted expression of *TP53I3* was spatially enriched in areas with architectural disruption or nuclear atypia (Table 3 and Supplementary Figure S4), suggesting that the model may detect regions undergoing genotoxic stress or p53 pathway activation, both of which have implications for tumor aggressiveness and treatment response in LUAD ^94^. Regions with high *TP53I3* expression localized to tumor nests with marked nuclear pleomorphism and structural disruption, consistent with stress-induced p53 signalling (Supplementary Figure S8-a) ^95^, while areas with low *TP53I3* expression showed preserved epithelial organization and uniform nuclei, appearing morphologically uniform and possibly indicative of lower DNA damage response activity (Supplementary Figure S8-b) ^96^.

On IHC images, TP53I3 demonstrated diffuse cytoplasmic staining with a granular pattern in tumor cells, with varying intensity in regions showing nuclear atypia and architectural disorganization (Figure 7). The tumor cells showcasing weaker expression matched a cluster of red-pixels on XpressO-lung’s heatmaps showcasing high probability (86%) for low-expression (Figure 7). While those showcasing higher expression matched to scattered cluster of yellow to cyan pixels showcasing low probability (14%) of high-expression (Figure 7 and Table 4). This variation in spatial expression of TP53I3 highlights the heterogeneity of tumor cells with respect to the p53 pathway activation and cellular stress response in lung adenocarcinoma (Figure 7).

**Figure 7.**
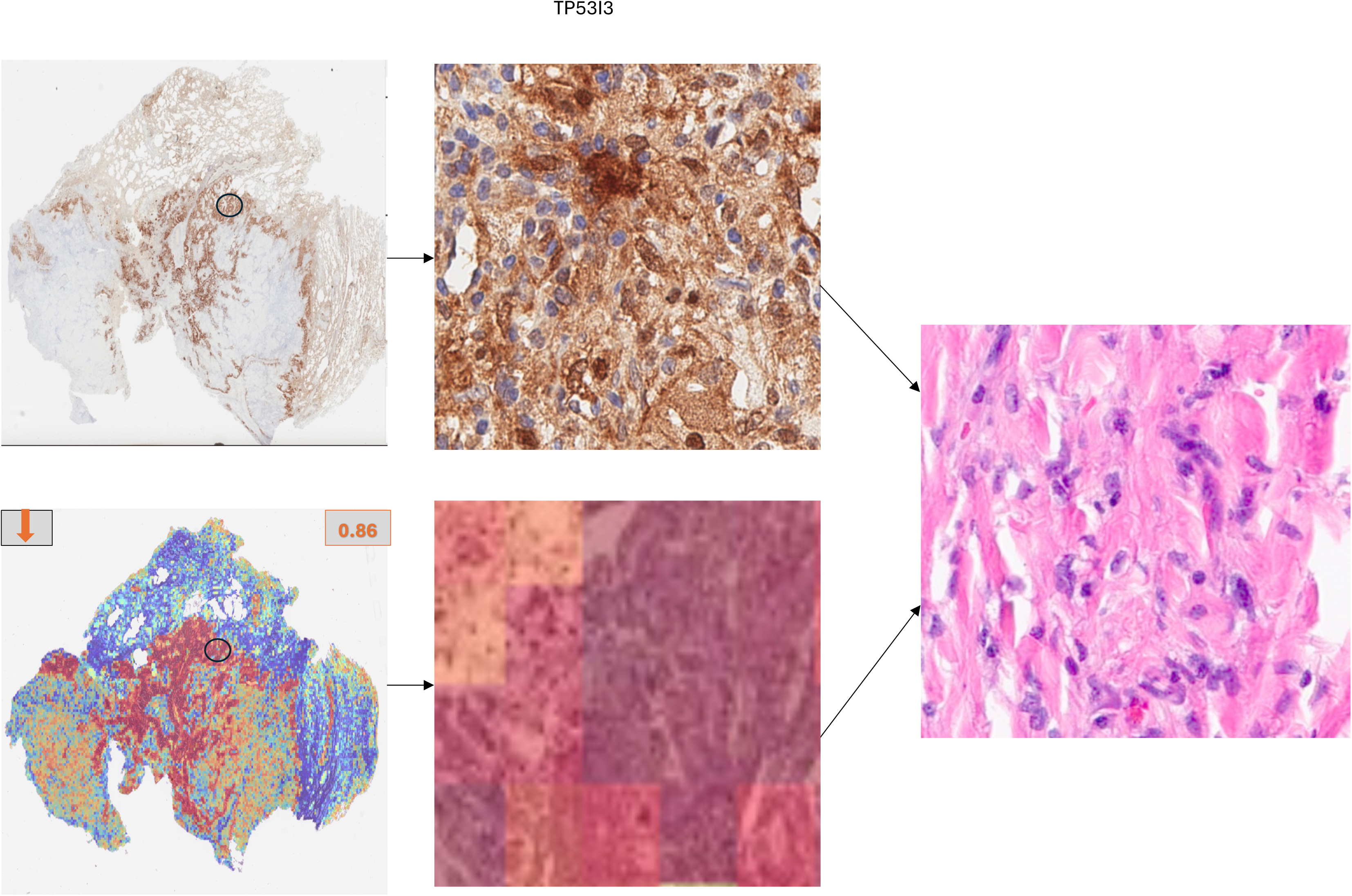
Immunohistochemistry (IHC), model attention maps and corresponding H&E views for TP53I3 expression in LUAD tissue sections. Representative cases showing spatial correspondence between IHC staining patterns and underlying histomorphology.

#### f FOXO1

FOXO1 is a tumor-suppressor transcription factor involved in cell cycle regulation and apoptosis, frequently inactivated in LUAD^83^. In the attention heatmaps, regions predicted to have low FOXO1 expression localized to tumor cell clusters with enlarged nuclei and irregular glandular architecture (Supplementary Figure S9). These features were consistently observed across high-attention regions, indicating that reduced FOXO1 expression is associated with morphologically aggressive tumor compartments characterized by nuclear atypia and architectural disorganization^83^.

On IHC images, FOXO1 demonstrated heterogeneous nuclear staining across tumor cells, with intermixed regions of higher expression, weaker expression and areas with minimal to absent staining (Figure 8). This spatial variability was reflected in the XpressO-Lung heatmaps, where clusters of red pixels corresponded to regions with higher expression probability (54%), while yellow to cyan pixels indicated regions with relatively lower expression probability (46%) (Figure 8 and Table 4). On H&E, these regions were characterized by architectural disorganization and nuclear atypia, including irregular nuclear contours and variability in nuclear size and chromatin pattern (Figure 8). The spatial overlap between these morphologic features and regions of variable FOXO1 expression suggests that the model may be capturing underlying tumor heterogeneity related to transcriptional regulation and cellular state differences in LUAD.

**Figure 8.**
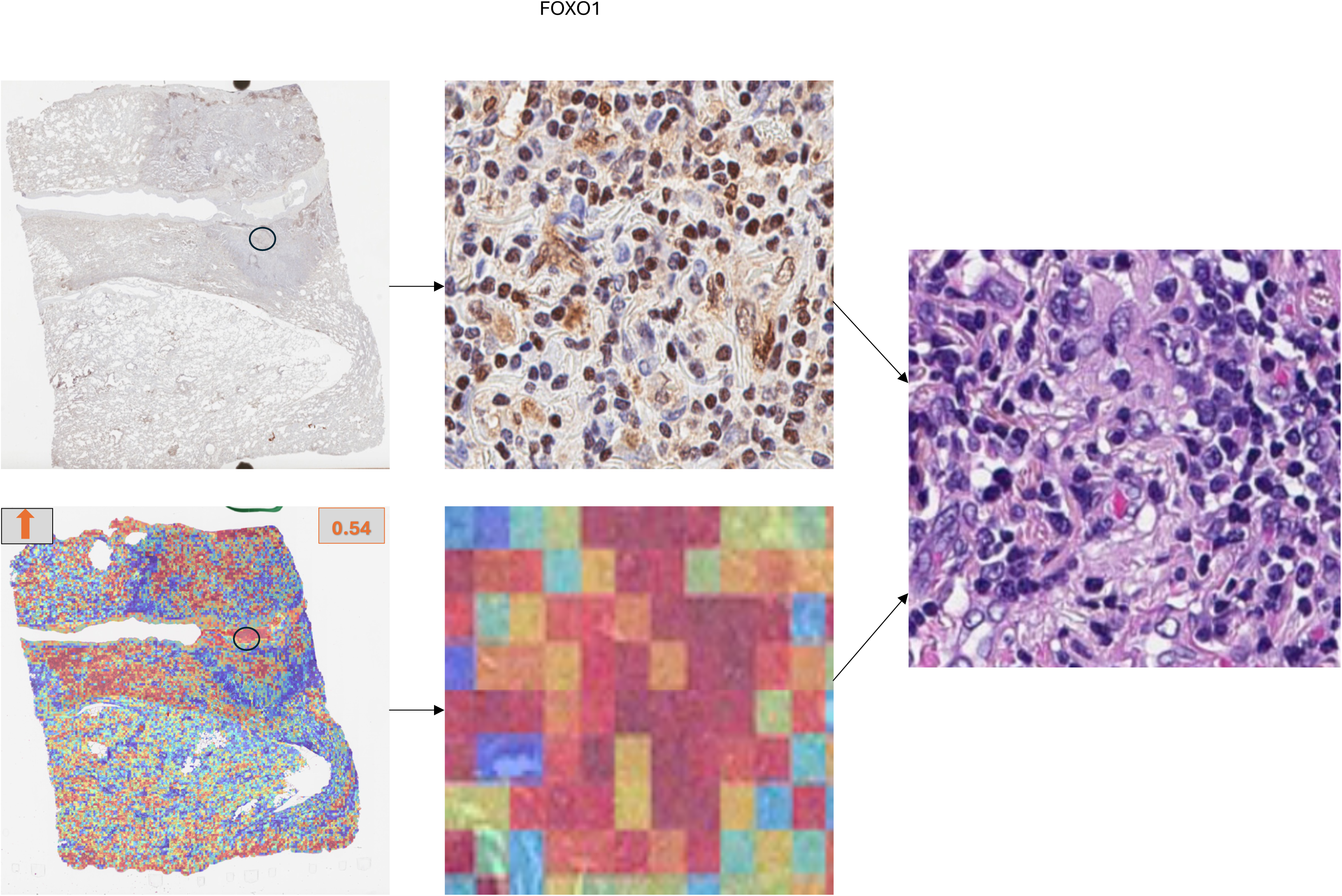
Immunohistochemistry (IHC), model attention maps and corresponding H&E views for FOXO1 expression in LUAD tissue sections. Representative cases showing spatial correspondence between IHC staining patterns and underlying histomorphology.

#### g KEAP1

KEAP1 is a tumor suppressor that negatively regulates the NRF2 oxidative stress pathway. Loss or inactivation of KEAP1 leads to persistent NRF2 activation, promoting tumor cell survival, metabolic adaptation, and therapy resistance in LUAD^108^. In our model, low KEAP1 expression was associated with attention focused on densely packed glandular regions composed of elongated nuclei (Supplementary Figure S10).

On IHC analysis, KEAP1 demonstrated cytoplasmic staining within tumor cells, with variable intensity across regions, ranging from moderate to strong expression (Figure 9). This spatial variability was reflected in the attention heatmaps, where clusters of red pixels corresponded to regions with higher expression probability (79%), while interspersed yellow to cyan pixels indicated regions with comparatively lower expression probability (21%) (Figure 9 and Table 4). On corresponding H&E sections, these regions were characterized by nuclear pleomorphism, atypical mitoses and multinucleated cells. The spatial concordance between KEAP1 expression patterns and these morphologic features suggests that the model may be capturing heterogeneity associated with oxidative stress response pathways and tumor progression in LUAD.

**Figure 9.**
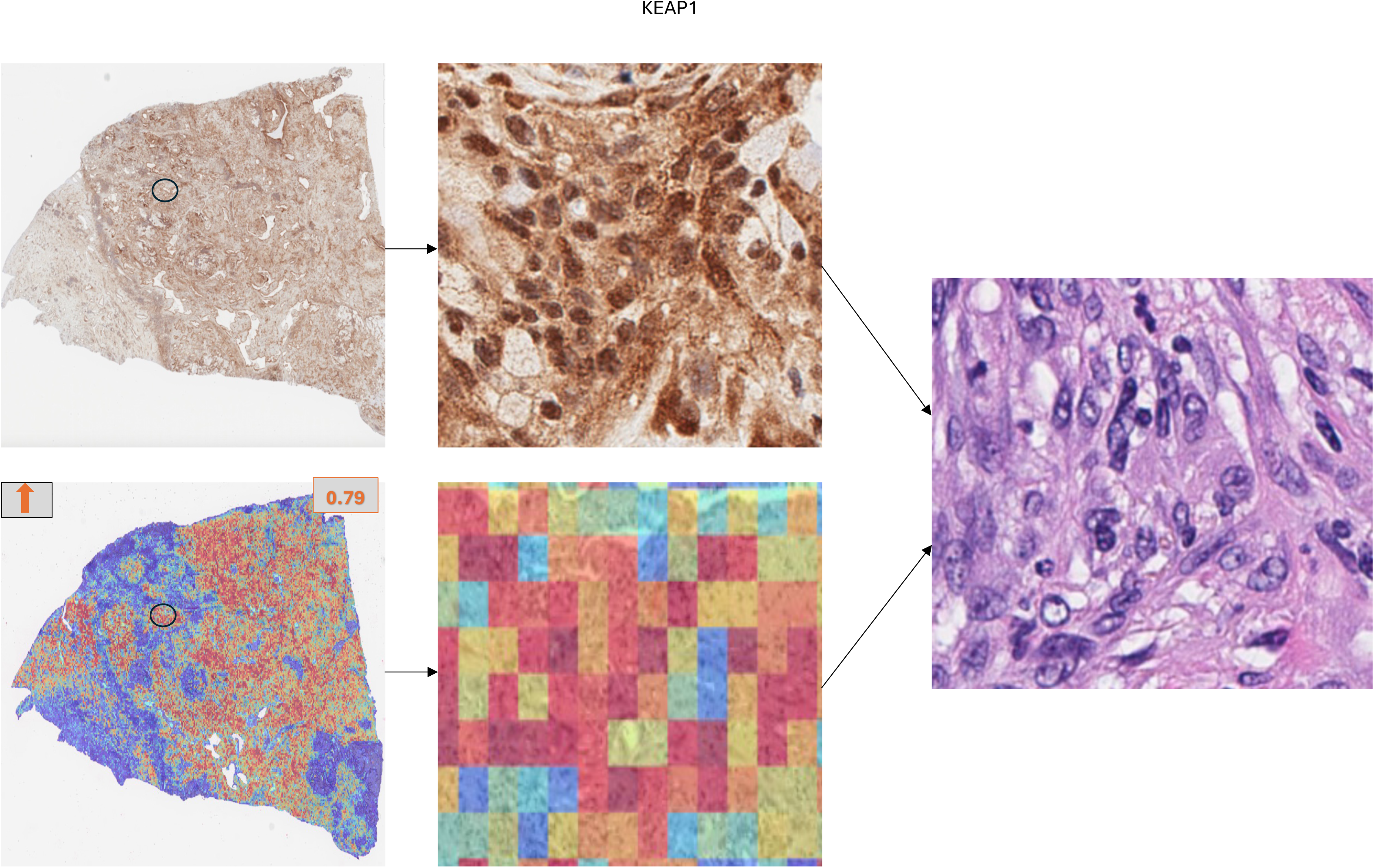
Immunohistochemistry (IHC), model attention maps and corresponding H&E views for KEAP1 expression in LUAD tissue sections. Representative cases showing spatial correspondence between IHC staining patterns and underlying histomorphology.

#### h RB1

RB1 is a tumor suppressor that regulates the G1/S cell cycle checkpoint and its loss leads to uncontrolled proliferation^109^. In our model, regions predicted as low RB1 expression showed attention on densely packed tumor cells with enlarged, hyperchromatic nuclei, increased nuclear crowding. These areas lacked well-spaced glandular architecture and instead displayed compact cellular arrangements with reduced structural organization, reflecting RB1-deficient tumor morphology (Supplementary Figure S11). On IHC, RB1 demonstrated predominantly weak to absent nuclear staining in tumor cells, with a large proportion of cells lacking detectable signal and only scattered cells showing focal nuclear positivity (Figure 10). This pattern is consistent with overall low expression. The XpressO-Lung heatmaps reflected this distribution, with regions of red pixels corresponding to higher probability of low expression (58%), and only some yellow to cyan pixels indicating regions with relatively higher expression probability (42%) (Figure 10 and Table 4). On corresponding H&E sections, these regions were characterized by irregular nuclei and variability in nuclear size, consistent with a poorly differentiated phenotype. The predominance of low RB1 expression across these morphologically atypical regions suggests that the model may be capturing tumor areas associated with altered cell cycle regulation and increased proliferative potential in LUAD.

**Figure 10.**
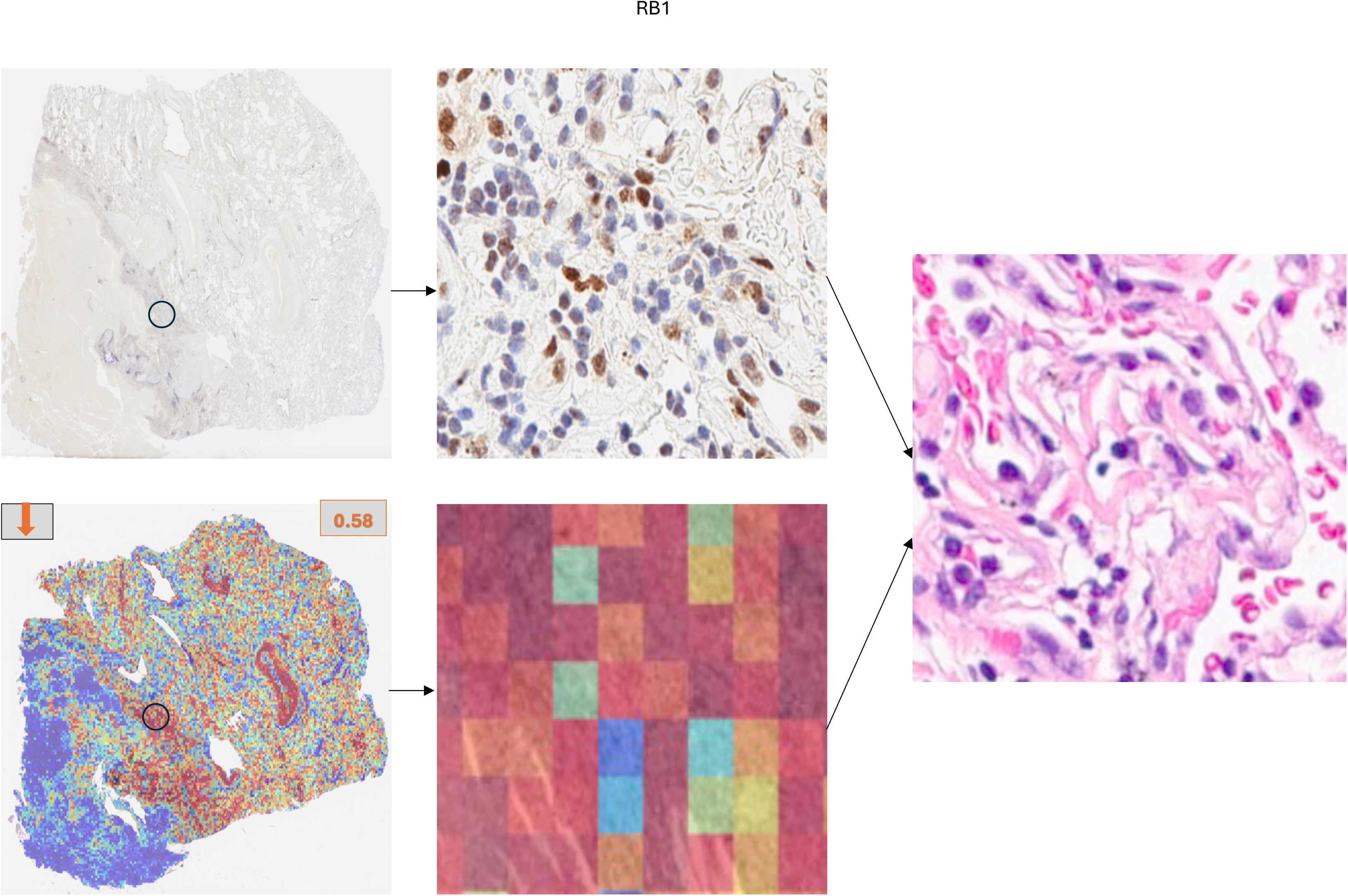
Immunohistochemistry (IHC), model attention maps and corresponding H&E views for RB1 expression in LUAD tissue sections. Representative cases showing spatial correspondence between IHC staining patterns and underlying histomorphology.

#### i TP53

TP53 is a tumor suppressor that regulates DNA damage response and cell cycle control and its alteration is associated with genomic instability and aggressive tumor behavior ^110^. In our model, high TP53 expression localized to regions with densely packed tumor cells showing enlarged, pleomorphic, hyperchromatic nuclei and marked nuclear crowding, reflecting proliferative morphology. In contrast, low TP53 expression localized to regions with compact tumor cell clusters exhibiting round nuclei, reduced pleomorphism, and less pronounced nuclear atypia, despite increased cellular density (Supplementary Figure S12). On immunohistochemical analysis, TP53 showed strong, diffuse nuclear staining in the majority of tumor cells, consistent with aberrant p53 accumulation and supporting a TP53-altered phenotype^110^ (Figure 11). In contrast, the heatmaps showed a heterogeneous mix of red, yellow, cyan, and blue pixels within the corresponding region, without a consistent spatial pattern aligning with the strong IHC signal (Figure 11 and Table 4). The lack of clear spatial concordance between the uniform IHC staining and heterogeneous model predictions suggests a discordance between predicted transcriptomic expression and protein-level expression, potentially reflecting differences in transcriptional and post-transcriptional regulation.

**Figure 11.**
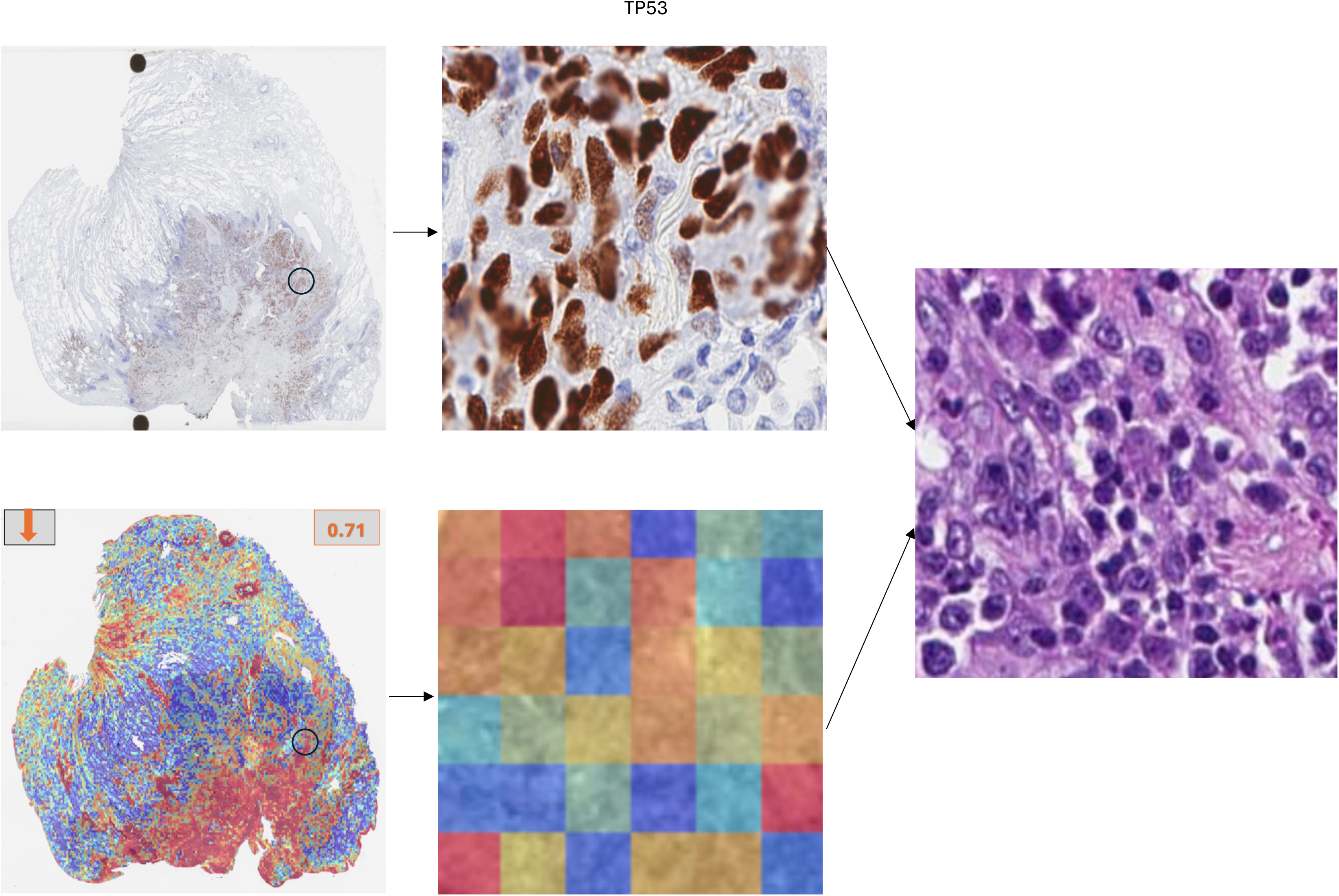
Immunohistochemistry (IHC), model attention maps and corresponding H&E views for TP53 expression in LUAD tissue sections. Representative cases showing spatial correspondence between IHC staining patterns and underlying histomorphology.

The biomarker pairs identified through our model also align with disease associations and pathways highlighted in STRING (version 12.5) ^111^, reinforcing their biological relevance. For instance, *CD8A*, *CDKN2A*, and *KRT7* showed strong associations with non-small cell lung carcinoma, vulvar carcinoma, and melanoma, with a very low false discovery rate in the Disease-gene Associations (DISEASES) ^112^ (Supplementary Figure 13-a). In this context, the FDR represents the probability that a reported association is a false positive after adjusting for multiple hypothesis testing, and values near zero indicate highly confident enrichment. Literature linked to these markers underscores their translational importance (Supplementary Figure S13-b): case reports highlight how immune contexture and PD-L1–driven dynamics in lung cancer can shape outcomes, implicating *CD8A*, *CDKN2A*, and *KRT7* in pathways that govern both therapeutic resistance and progression across subtypes ^113^. Other studies emphasize their roles in intratumoral heterogeneity that drives drug resistance ^114^ and spatial heterogeneity influencing survival in lung adenocarcinoma patients ^115^. Furthermore, guidelines for diagnostic immunohistochemistry in lung cancer recommend markers like *KRT7* and *CDKN2A* as lineage classifiers, directly supporting their inclusion in predictive modelling ^116^.

Additionally, STRING associations among *TTF1*, *KRT7*, *CDKN2A*, and *NAPSA* connect these canonical lung adenocarcinoma markers to both disease associations (NSCLC, intestinal cancer, vulvar carcinoma) and specific pathways such as surfactant metabolism, in which *TTF1* and *NAPSA* play key regulatory roles (Supplementary Figure S14-a). Here again, FDR-adjusted enrichment values were very low, indicating robust statistical confidence. Literature linked to this cluster highlights their clinical importance in molecularly complex or therapy-resistant settings (Supplementary Figure S14-b): ALK-rearranged lung tumors lacking typical lineage markers ^120^, ALK fusion–driven resistance to osimertinib ^121^, and histomorphological transformation from NSCLC to small cell carcinoma after immunotherapy ^122^ all emphasize the diagnostic and prognostic value of these markers. Meta-analyses distinguishing adenocarcinoma from squamous carcinoma further reinforce the utility of *KRT7* and *TTF1* as diagnostic anchors ^123^. Notably, STRING highlights a local cluster of *NAPSA* and *KRT7* with signet-ring cell adenocarcinoma, a rarer subtype, while in our LUAD cohort, these markers instead aligned with classical adenocarcinoma morphologies, reinforcing their broader lineage-defining role. (Supplementary Figure S14-a).

These STRING-derived connections illustrate how the biomarker pairs identified by our model intersect with established cancer pathways, disease associations, and prior studies.

## Discussion

In this study, we present XpressO-Lung, an explainable deep learning model that projects gene expression heterogeneity spatially in LUAD tumors and their microenvironment on H&E-stained Dx-WSIs using their corresponding bulk-transcriptomics data. The model was validated and interpreted using immunohistochemistry on an external set of clinical samples at DH and was found to capture histomorphologic patterns predictive of both immune and epithelial gene expression signatures within LUAD tissues. Our results demonstrate that the model not only recapitulates known LUAD biomarkers, i.e., *NAPSA*, *TTF1*, *KRT7*, *CDKN2A*, *CD8A*, *TP53I3*, FOXO1, *KEAP1, RB1* and *TP53* but also reveals their biologically meaningful spatial expression patterns either in a paired or individual format, mirroring underlying tumor and TME interactions.

Across these biomarkers, the model’s predictive performance ranged from AUC 0.64-0.92, providing a strong foundation for interpreting the spatial patterns it uncovered across distinct epithelial lineages, proliferative states, and immune–stromal niches in LUAD.

XpressO-Lung uncovered diverse epithelial lineages and proliferative states within LUAD. NAPSA and TTF1 frequently co-localized in cohesive glandular nests, anchoring well-differentiated adenocarcinoma clones ^83–85^. In contrast, KRT7-bright/CDKN2A-silent regions marked 9p21-loss proliferative clones ^87–88^, and NAPSA-bright/CDKN2A-silent zones reflected alveolar-differentiated yet cell-cycle-unrestrained phenotypes, both linked to poor prognosis yet potentially targetable by CDK4/6 or PRMT5 inhibitors ^89–90^. These spatial phenotypes align with STRING associations connecting NAPSA, TTF1, KRT7 and CDKN2A to surfactant metabolism, cell-cycle regulation, and NSCLC pathways ^116–119^. Beyond epithelial compartments, the model also mapped immune and stromal microenvironments with clear spatial resolution. CD8A frequently rimmed KRT7-positive tumor islands, forming lymphocytic cuffs characteristic of an immune-inflamed TME ^86^.

XpressO-Lung also identified stress-responsive and therapy-relevant niches within LUAD tissue. TP53I3 marked disordered regions with nuclear pleomorphism, hinting at p53 pathway activation and oxidative stress, while its low expression was observed in morphologically uniform zones ^95^. Extending these findings, canonical tumor suppressors further refined these spatial states: low FOXO1 expression localized to gland-forming regions with irregular architecture and nuclear atypia, indicating disrupted epithelial organization despite retained differentiation^83^; low KEAP1 expression highlighted densely packed glandular regions with elongated nuclei, consistent with oxidative stress-adapted tumor compartments ^108^; and low RB1 expression marked highly cellular, poorly organized regions with nuclear crowding and hyperchromasia, reflecting unchecked proliferation ^109^. In parallel, TP53 alterations captured a spectrum of nuclear atypia, with high TP53 expression associated with pleomorphic, densely packed tumor regions and low TP53 expression linked to more uniform but compact cellular clusters ^110^. Notably, these morphologic patterns showed concordance with immunohistochemical findings on an external dataset, reinforcing that model-predicted spatial features reflect underlying tumor biology. Together, these patterns illustrate how XpressO-Lung characterizes the spatial architecture of the LUAD tumor-TME interface, revealing how morphology encodes both lineage state and treatment-relevant phenotypes.

While XpressO-Lung demonstrates strong predictive performance and spatial interpretability, certain limitations remain. Although the external validation cohort was limited to eight tissue samples, further validation of results was performed using IHC, a standard-of-care technique that provides direct protein-level assessment and serves as a robust orthogonal modality to transcriptomic predictions. This enables meaningful biological validation through spatial concordance between model-predicted gene expression contributing regions and IHC (protein expression) staining patterns, even in smaller cohorts. Additionally, this approach offers a cost-effective and practical strategy, particularly in funding-constrained or resource-limited settings where large, multi-modal datasets may not be readily available. Future work will focus on expanding validation across larger, independent, and multi-institutional cohorts to further establish the generalizability of the model. In addition, although the model is trained to associate bulk expression with local histomorphologic features and validated orthogonally with spatial IHC patterns, a secondary validation at the subcellular level would be helpful in near future. In future work, we plan to close this gap by performing targeted subcellular spatial transcriptomics and proteomics experiments on the same in-house LUAD tissues. This will enable finer-grained validation of XpressO-Lung’s attention maps and expression predictions, further strengthening its clinical relevance and utility in translational settings.

XpressO-Lung advances the field by offering an accessible, scalable, and interpretable tool for morpho-genomic profiling in LUAD. It bridges WSI, bulk-transcriptomics and IHC using standard clinical slides, circumventing the need for expensive molecular assays, and enabling broader biomarker discovery, stratification of immune versus epithelial tumor states, and improved patient-specific therapy selection. In deep learning for computational pathology, interpreting model outputs is as critical as achieving high predictive performance and XpressO-Lung delivers on both fronts. The model’s framework reliably infers spatial gene expression patterns from histologic images and highlights biologically meaningful tumor-TME interactions. The model was trained, validated and tested using a carefully structured design that ensured sufficient degrees of freedom for each gene of interest, and incorporated a custom script for gene expression analysis to systematically reveal morphology-gene expression relationships and then validate it further with ground truth IHC based protein expression patterns. By combining rigorous methodology with transparent, interpretable outputs and external ground truth validation, XpressO-Lung offers a powerful complement to molecular profiling, with the potential to inform clinical decision-making, advance biomarker discovery, and support precision medicine in LUAD, particularly in resource-limited settings.

## Acknowledgements

The authors acknowledge the support of the Pathology Shared Resource (RRID: SCR_023479) at the Dartmouth Cancer Center with NCI Cancer Center Support Grant P30 CA023108, for the administration of Immunohistochemistry (IHC) experiment.

## SUPPLEMENTARY FIGURE LEGENDS

**Supplementary Figure S1.** Attention heatmaps and corresponding H&E views for *NAPSA* and *TTF1* expression predictions in LUAD whole-slide images. (a) Case (TCGA-55-A493) showing low *NAPSA* expression (a-i) and high *TTF1* expression (a-ii). (b) Case (TCGA-75-7027) showing low expression for both *NAPSA* (b-i) and *TTF1* (b-ii). High or low expression status is indicated by red arrows, with the predicted probability of high or low expression (*p_0*) shown in boxes. Black circles mark regions of interest selected for zoom-in visualization.

**Supplementary Figure S2.** Attention heatmaps and corresponding H&E views for *NAPSA* and *TTF1* expression predictions in LUAD whole-slide images. (a) Case (TCGA-55-1592) showing low *NAPSA* expression (a-i) and high *TTF1* expression (a-ii). High or low expression status is indicated by red arrows, with the predicted probability of high or low expression (*p_0*) shown in boxes. Black circles mark regions of interest selected for zoom-in visualization.

**Supplementary Figure S3.** Attention heatmaps and corresponding H&E views for *CD8A* and *KRT7* expression predictions in LUAD whole-slide images. (a) Case (TCGA-49-AAR3) showing high *CD8A* expression (a-i) and low *KRT7* expression (a-ii). (b) Case (TCGA-69-7973) showing high expression for both *CD8A* (b-i) and *KRT7* (b-ii). High or low expression status is indicated by red arrows, with the predicted probability of high or low expression (*p_0*) shown in boxes. Black circles mark regions of interest selected for zoom-in visualization. Areas labeled “T” indicate tumor nests, while areas labeled “TME” denote the surrounding tumor microenvironment.

**Supplementary Figure S4.** Attention heatmaps and corresponding H&E views for *CD8A* and *KRT7* expression predictions in LUAD whole-slide images. (a) Case (TCGA-44-5643) showing high *CD8A* expression (a-i) and high *KRT7* expression (a-ii). High or low expression status is indicated by red arrows, with the predicted probability of high or low expression (*p_0*) shown in boxes. Black circles mark regions of interest selected for zoom-in visualization. Areas labeled “T” indicate tumor nests, while areas labeled “TME” denote the surrounding tumor microenvironment.

**Supplementary Figure S5.** Attention heatmaps and corresponding H&E views for *KRT7* and *CDKN2A* expression predictions in LUAD whole-slide images. (a) Case (TCGA-44-8119) showing high *KRT7* expression (a-i) and low *CDKN2A* expression (a-ii). (b) Case (TCGA-97-A4M5) showing high *KRT7* expression (b-i) and low *CDKN2A* expression (b-ii). High or low expression status is indicated by red arrows, with the predicted probability of high or low expression (*p_0*) shown in boxes. Black circles mark regions of interest selected for zoom-in visualization.

**Supplementary Figure S6.** Attention heatmaps and corresponding H&E views for *NAPSA* and *CDKN2A* expression predictions in LUAD whole-slide images. (a) Case (TCGA-97-A4M1) showing high *NAPSA* expression (a-i) and low *CDKN2A* expression (a-ii). (b) Case (TCGA-55-7907) showing high *NAPSA* expression (b-i) and low *CDKN2A* expression (b-ii). High or low expression status is indicated by red arrows, with the predicted probability of high or low expression (*p_0*) shown in boxes. Black circles mark regions of interest selected for zoom-in visualization.

**Supplementary Figure S7.** Attention heatmaps and corresponding H&E views for *NAPSA* and *CDKN2A* expression predictions in LUAD whole-slide images. (a) Case (TCGA-55-8506) showing low *NAPSA* expression (a-i) and low *CDKN2A* expression (a-ii). (b) Case (TCGA-97-A4M5) showing high *NAPSA* expression (b-i) and low *CDKN2A* expression (b-ii). High or low expression status is indicated by red arrows, with the predicted probability of high or low expression (*p_0*) shown in boxes. Black circles mark regions of interest selected for zoom-in visualization.

**Supplementary Figure S8.** Attention heatmaps and corresponding H&E views for *TP53I3* expression predictions in LUAD whole-slide images. (a) Case (TCGA-05-4397) showing high *TP53I3* expression. (b) Case (TCGA-78-7153) showing low *TP53I3* expression High or low expression status is indicated by red arrows, with the predicted probability of high or low expression (*p_0*) shown in boxes. Black circles mark regions of interest selected for zoom-in visualization.

**Supplementary Figure S9. Attention heatmaps and corresponding H&E views for FOXO1 expression predictions in LUAD whole-slide images.** (a) Case (TCGA-MP-A4SY) showing low FOXO1 expression. (b) Case (TCGA-97-7554) showing low FOXO1 expression. Low expression status is indicated by red arrows, with the predicted probability of low expression (p) shown in boxes. Black circles mark regions of interest selected for zoom-in visualization.

**Supplementary Figure S10. Attention heatmaps and corresponding H&E views for KEAP1 expression predictions in LUAD whole-slide images.** (a) Case (TCGA-05-4430) showing low KEAP1 expression. (b) Case (TCGA-55-7913) showing low KEAP1 expression. Low expression status is indicated by red arrows, with the predicted probability of low expression (p) shown in boxes. Black circles mark regions of interest selected for zoom-in visualization.

**Supplementary Figure S11. Attention heatmaps and corresponding H&E views for RB1 expression predictions in LUAD whole-slide images.** (a) Case (TCGA-78-7540) showing low RB1 expression. (b) Case (TCGA-97-7547) showing low RB1 expression. Low expression status is indicated by red arrows, with the predicted probability of low expression (p) shown in boxes. Black circles mark regions of interest selected for zoom-in visualization.

**Supplementary Figure S12. Attention heatmaps and corresponding H&E views for TP53 expression predictions in LUAD whole-slide images.** (a) Case (TCGA-49-AAR2) showing low TP53 expression. (b) Case (TCGA-62-A46S) showing low TP53 expression. Low expression status is indicated by red arrows, with the predicted probability of low expression (p) shown in boxes. Black circles mark regions of interest selected for zoom-in visualization.

**Supplementary Figure S13.** (a) STRING disease–gene associations for *CD8A*, *CDKN2A*, and *KRT7* from DISEASES. (b) Reference publications linked to these markers. Node colors reflect supporting evidence categories; edges (green line) represent text-mining associations.

**Supplementary Figure S14.** (a) STRING Reactome pathway, network cluster, and disease gene associations for *TTF1*, *KRT7*, *CDKN2A*, and *NAPSA*. (b) Reference publications linked to this cluster. Node colors reflect supporting evidence categories; edges (green line) represent text-mining associations.

