## Supplementary Figures for "Interpreting and Validating a Deep Learning Model Predictive of Spatial Morphologic-Molecular Patterns in Lung Adenocarcinoma, Using Ground Truth Immunohistochemistry Images"

**NAPSA and TTF1**

**A-i)**

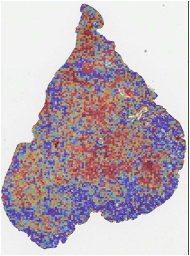

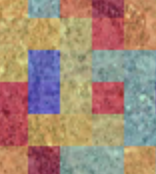

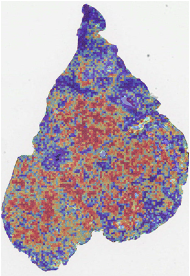

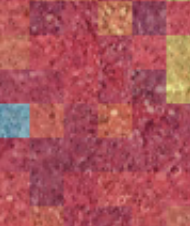

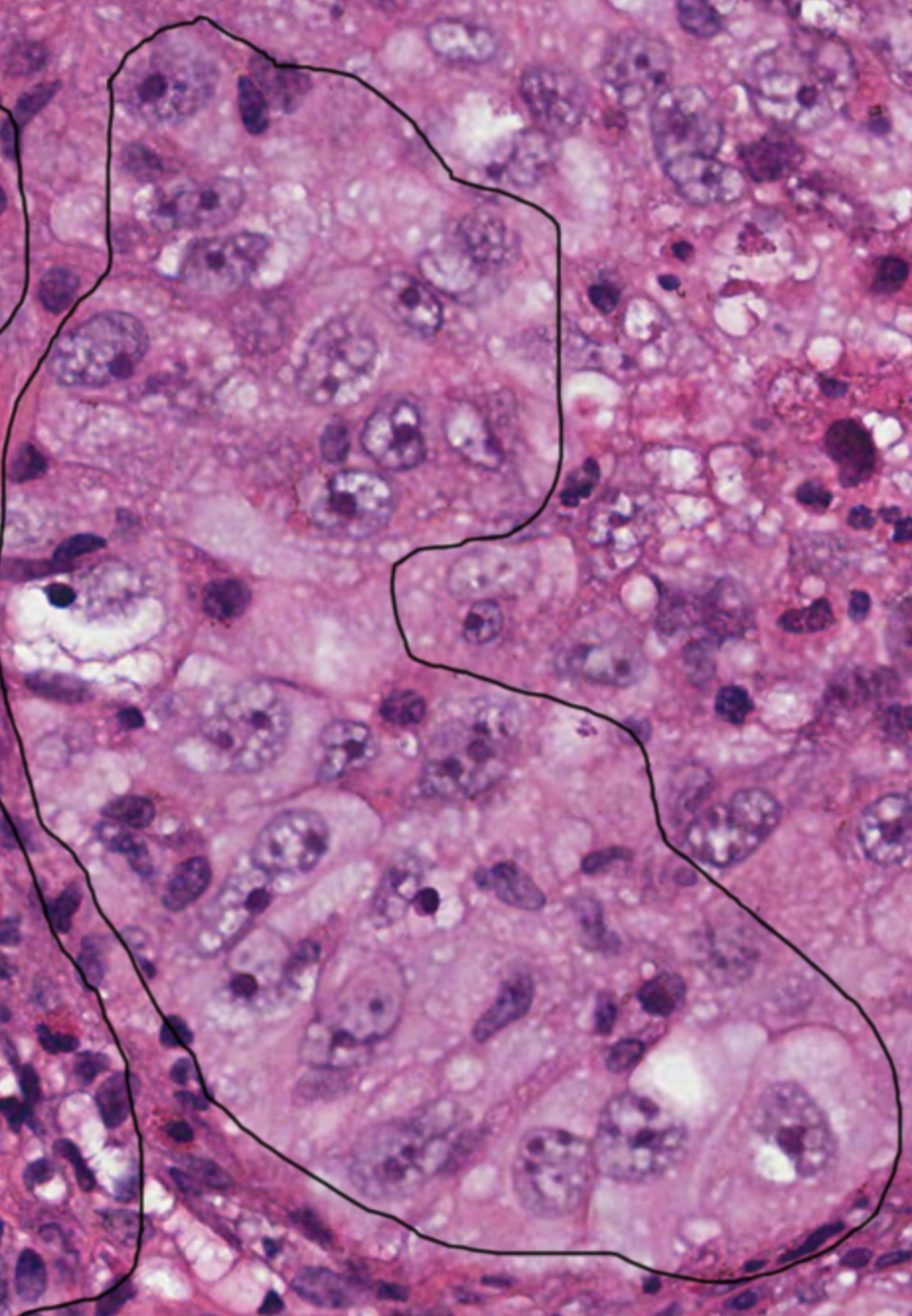

**0.61**

TTF1

NAPSA

**0.73**

**A-ii)**

**0.62**

**B-i)**

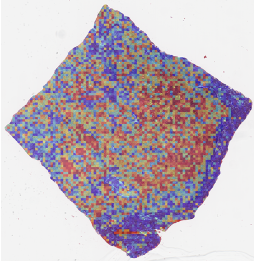

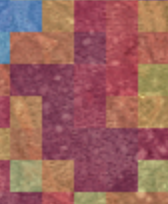

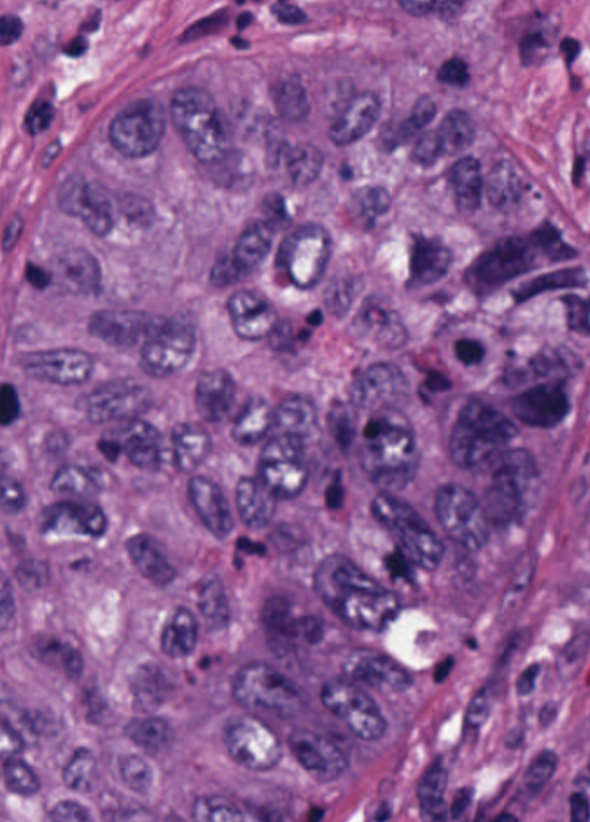

NAPSA

**B-ii)**

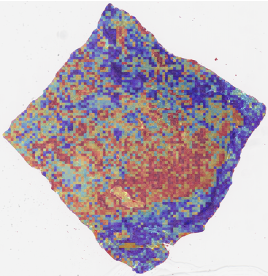

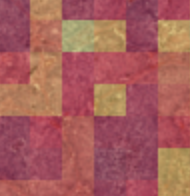

**0.54**

TTF1

**Supplementary Figure S1**

**NAPSA and TTF1**

**A-i)**

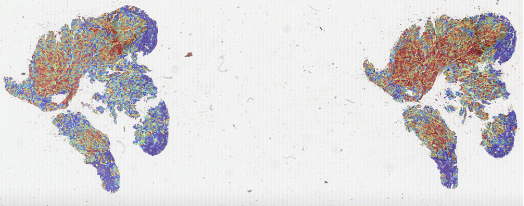

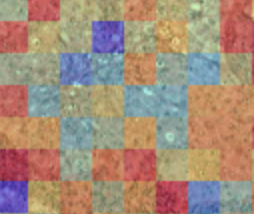

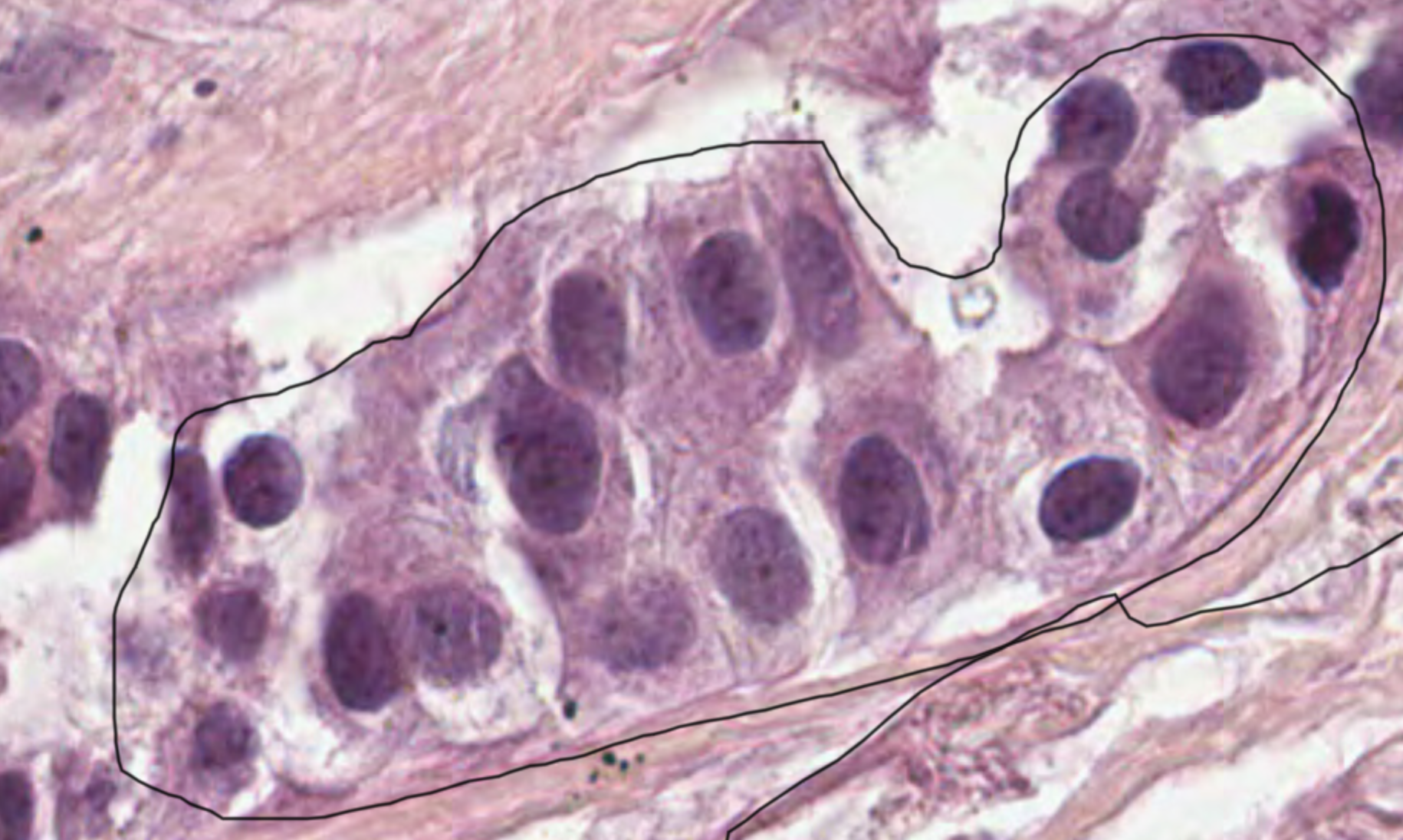

**0.73**

NAPSA

**0.80**

**A-ii)**

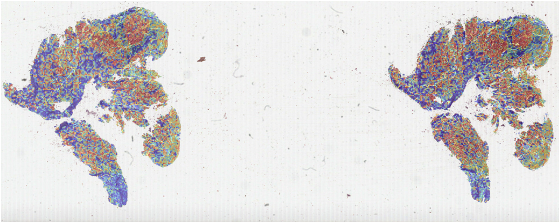

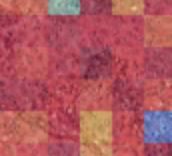

TTF1

**Supplementary Figure S2**

**CD8A and KRT7**

TME

**A-i)**

**0.61**

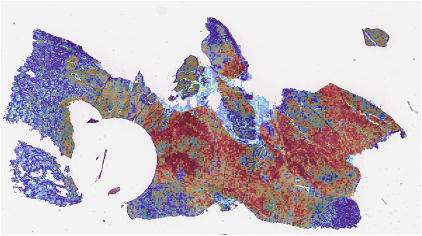

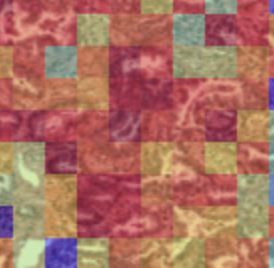

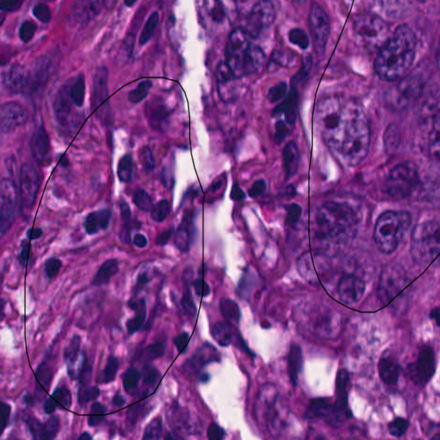

T

TME

**A-ii)**

CD8A

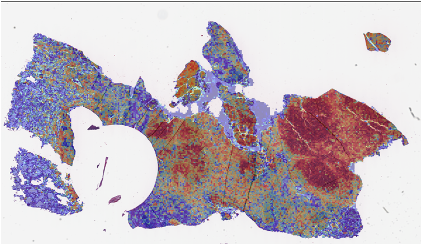

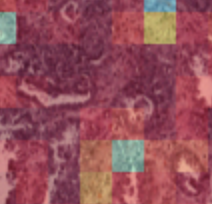

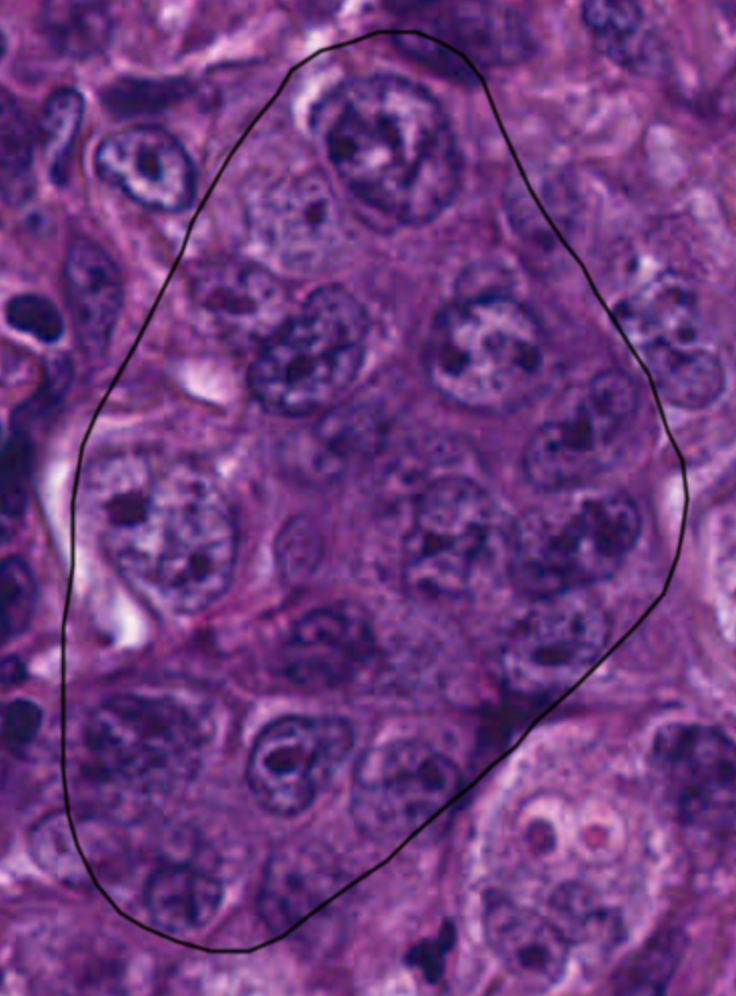

T

T

**0.69**

KRT7

TME

**B-i)**

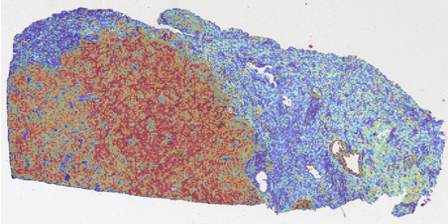

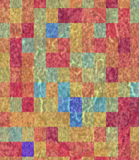

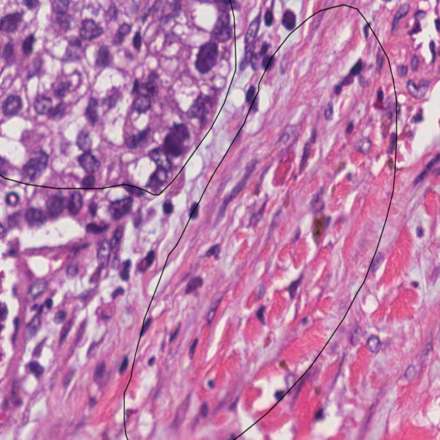

T

T

TME

**B-ii)**

**0.78**

**0.82**

CD8A

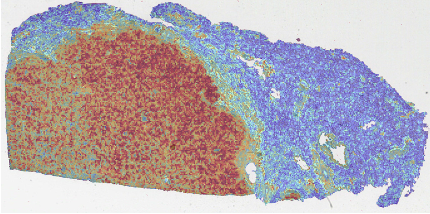

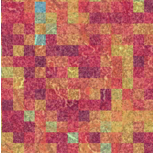

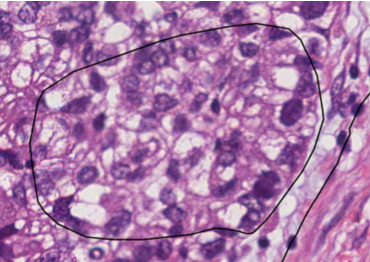

T

KRT7

**Supplementary Figure S3**

**CD8A and KRT7**

**A-i)**

TME

**0.88**

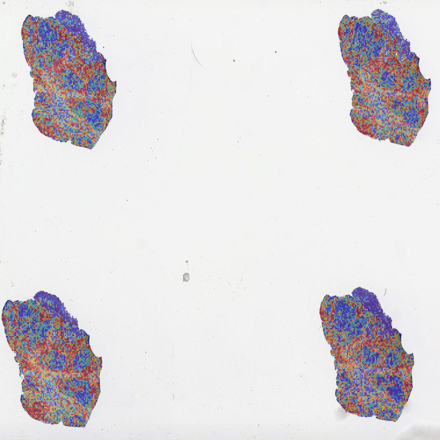

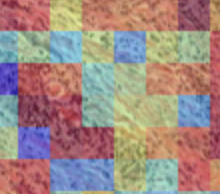

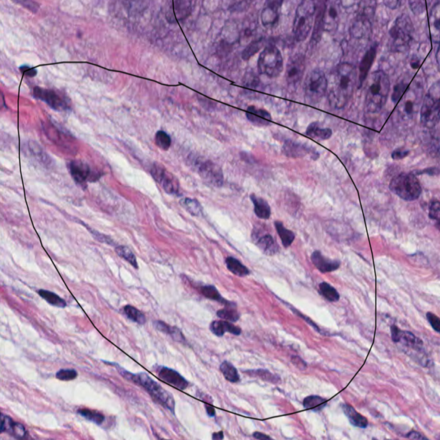

**A-ii)**

TME

T

CD8A

T

**0.74**

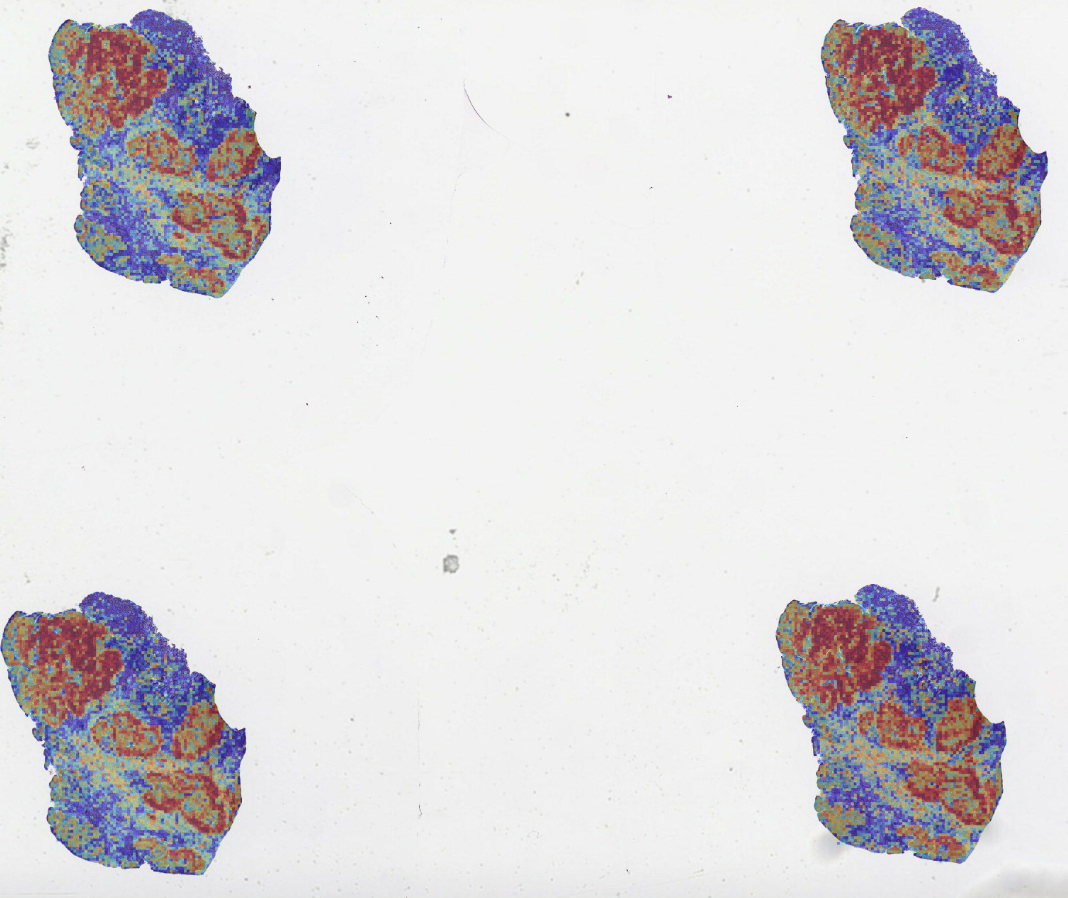

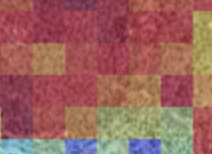

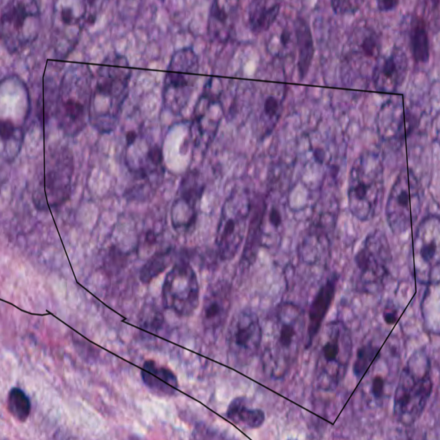

T

KRT7

**Supplementary Figure S4**

**KRT7 and CDKN2A**

**0.53**

**A-i)**

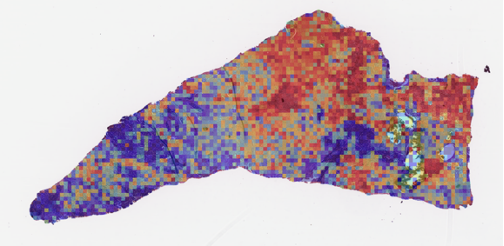

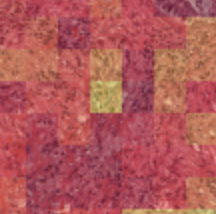

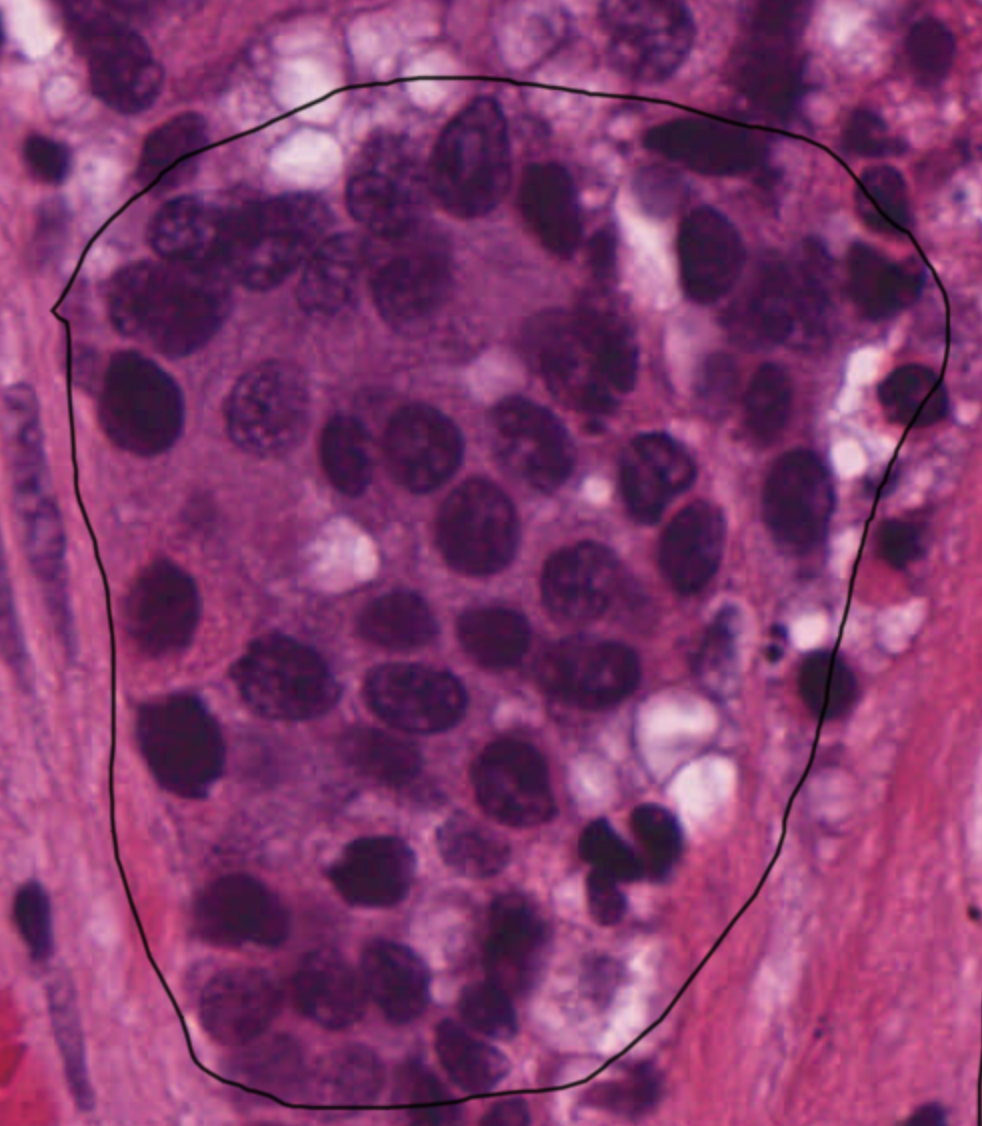

KRT7

**A-ii)**

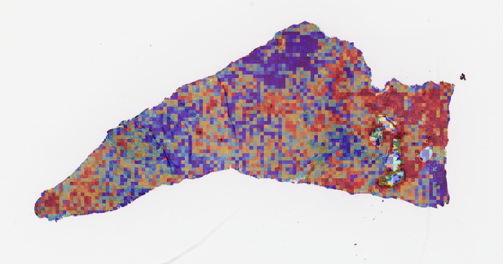

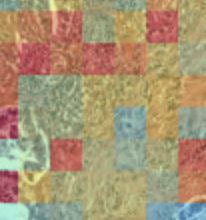

**0.76**

CDKN2A

**0.77**

**B-i)**

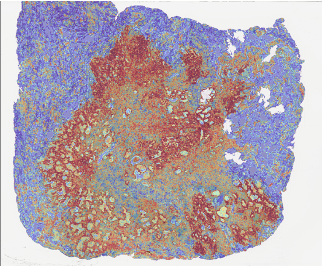

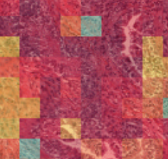

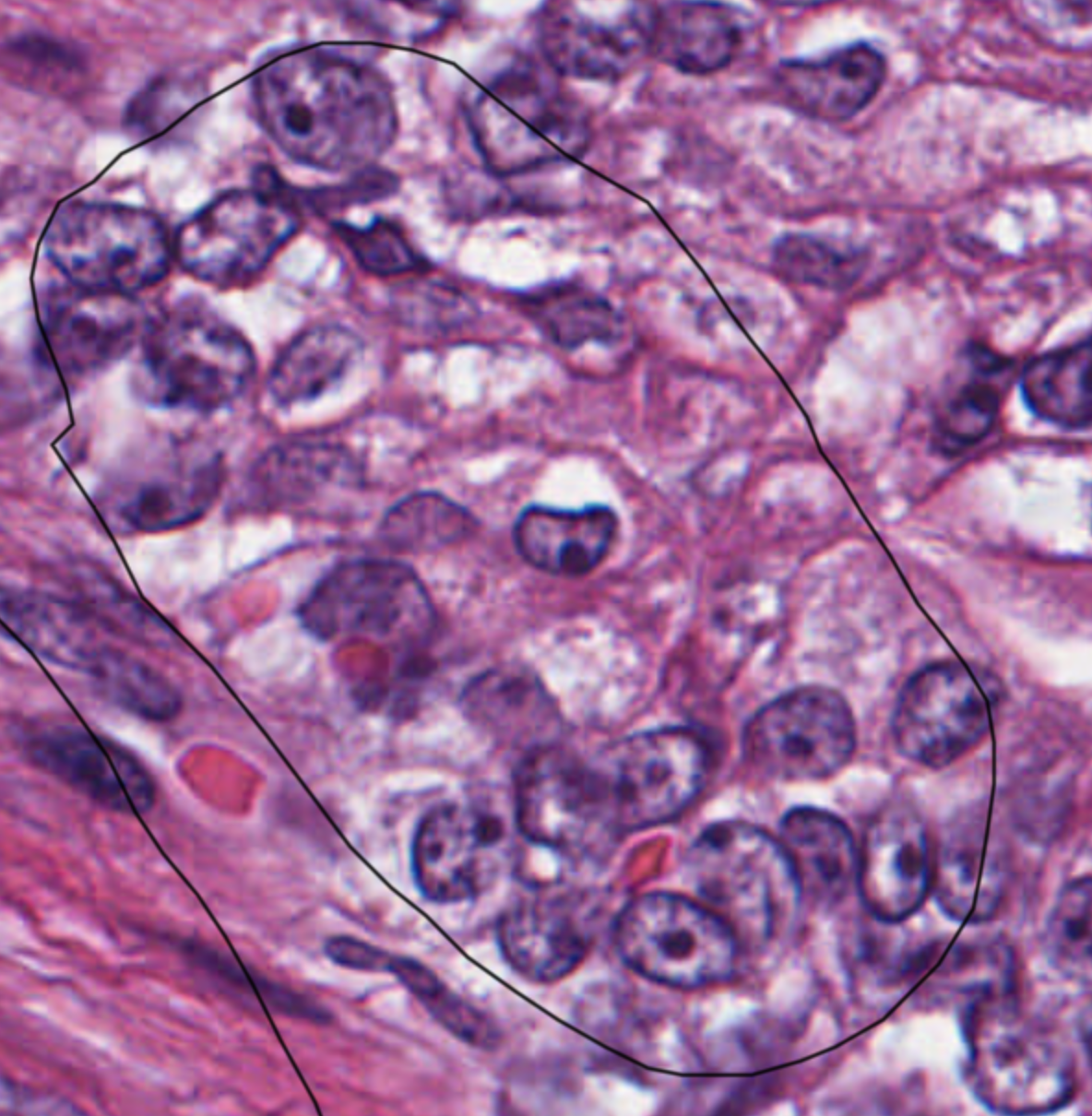

**0.66**

KRT7

**B-ii)**

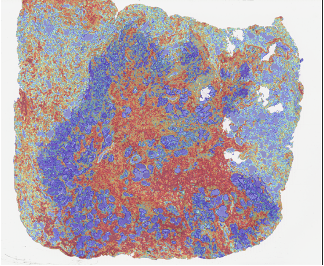

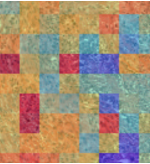

CDKN2A

**Supplementary Figure S5**

**NAPSA and CDKN2A**

**0.90**

**A-i)**

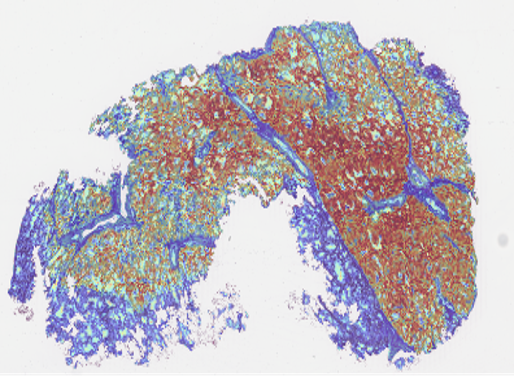

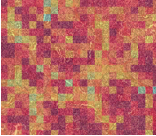

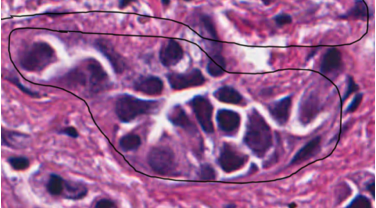

NAPSA

**A-ii)**

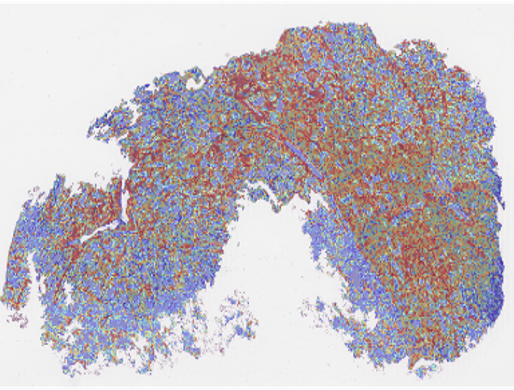

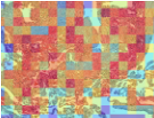

CDKN2A

**0.75**

**B-i)**

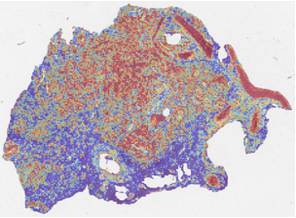

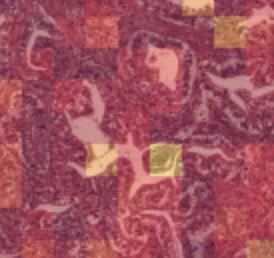

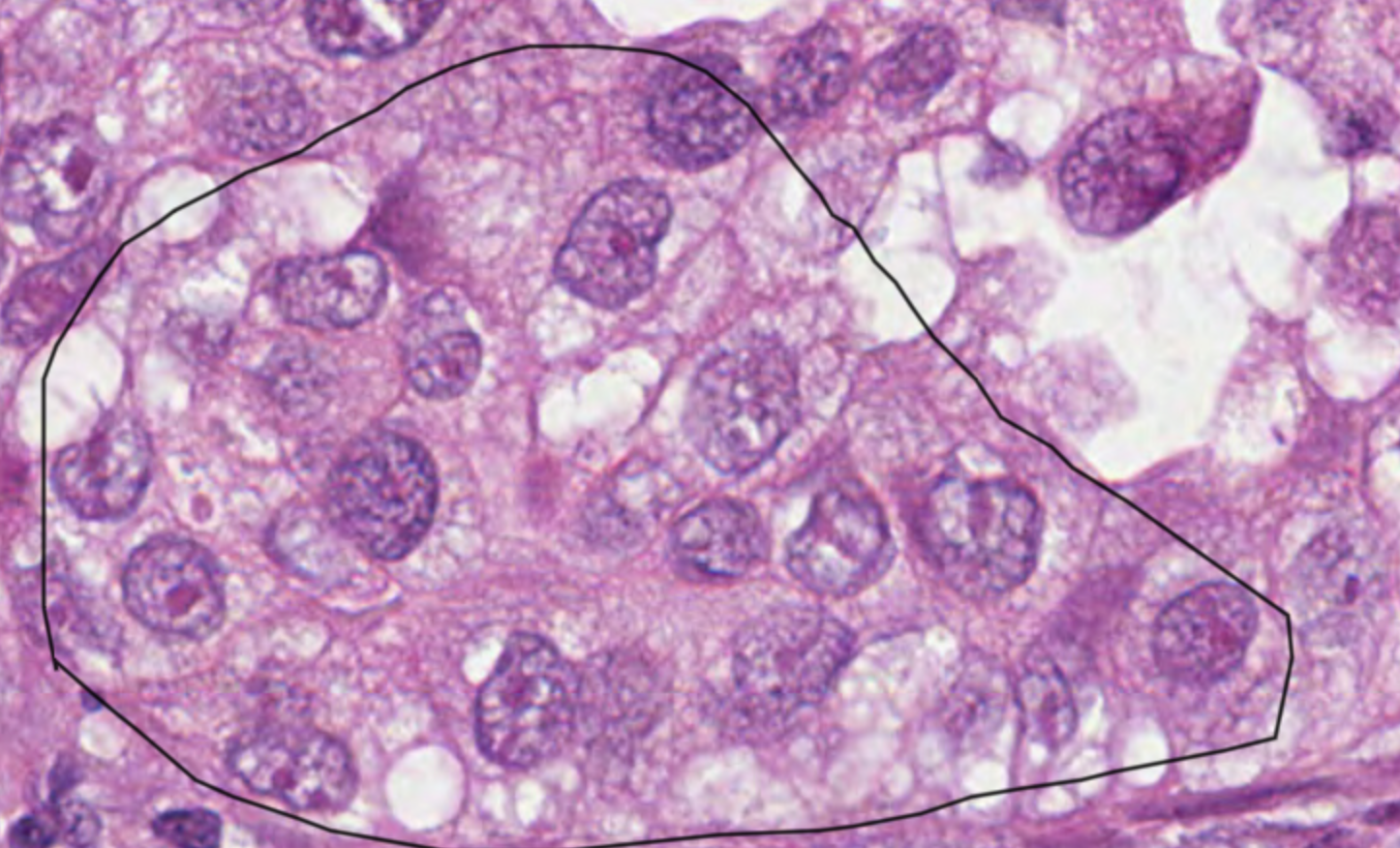

NAPSA

**0.54**

**B-ii)**

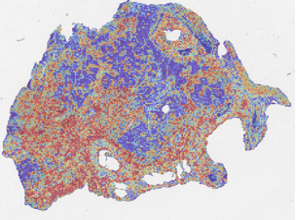

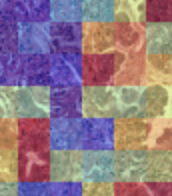

CDKN2A

**0.55**

**Supplementary Figure S6**

**NAPSA and CDKN2A**

**0.79**

**A-i)**

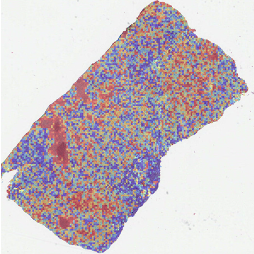

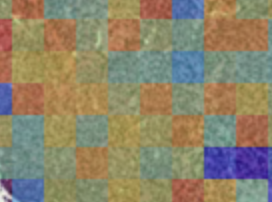

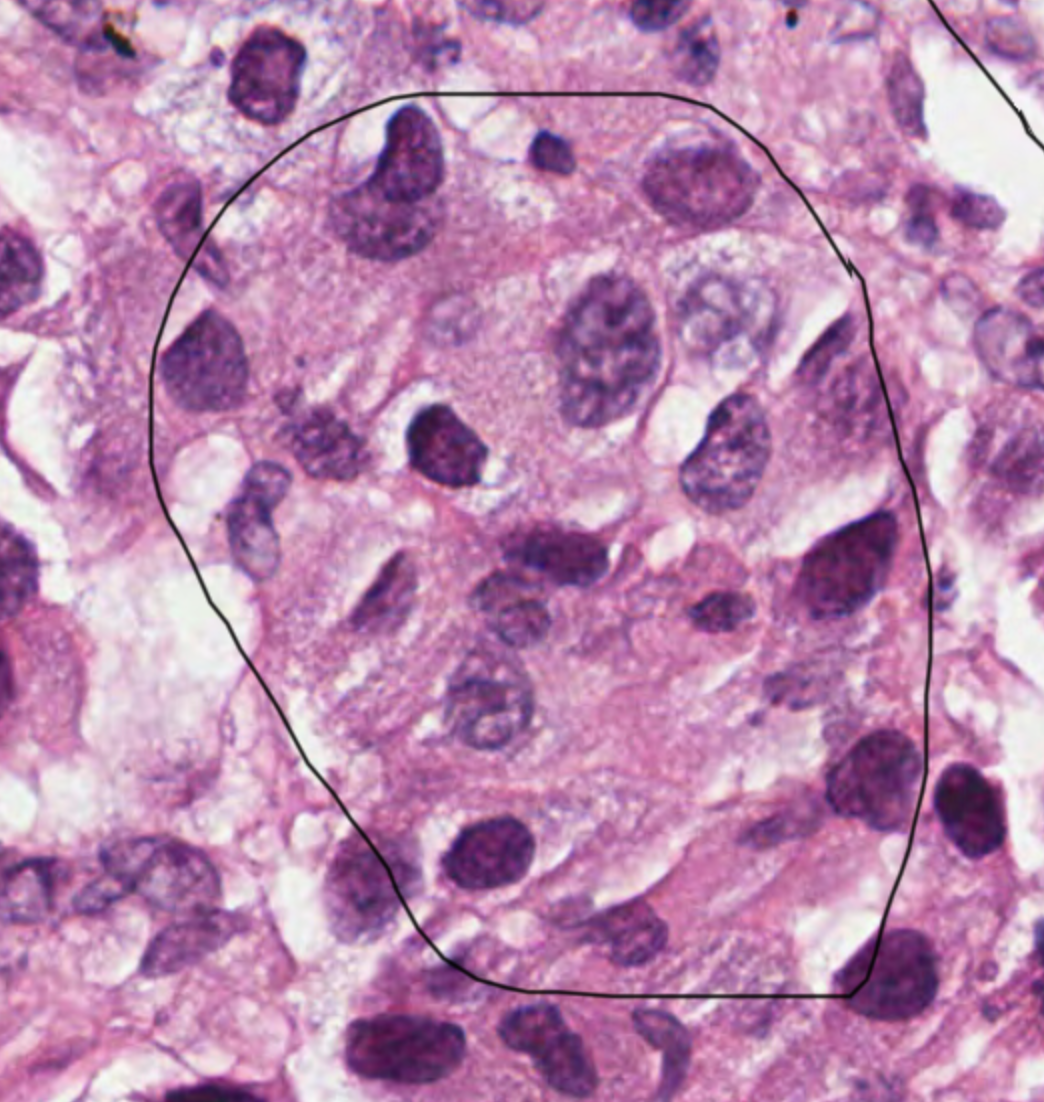

NAPSA

**A-ii)**

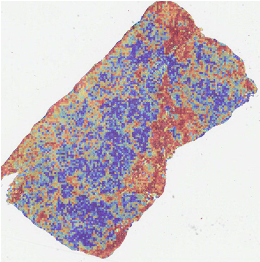

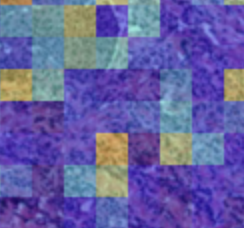

**0.63**

CDKN2A

**B-i)**

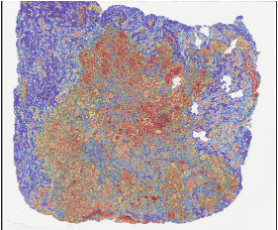

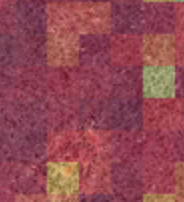

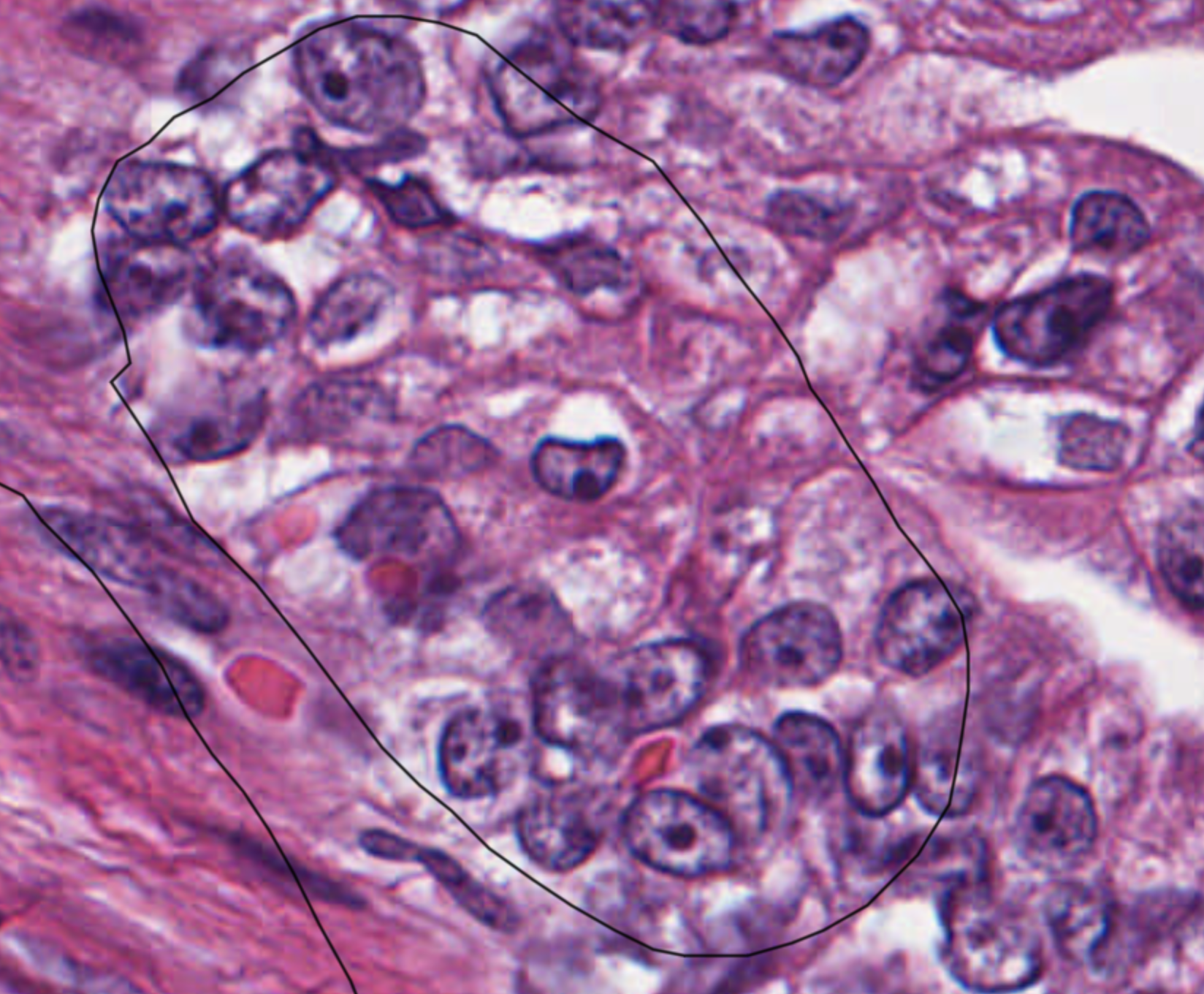

**0.52**

NAPSA

**B-ii)**

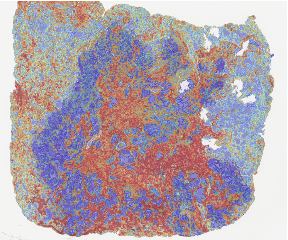

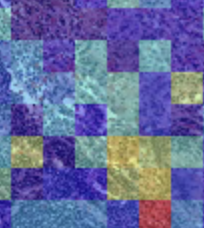

**0.66**

CDKN2A

**Supplementary Figure S7**

**TP53I3**

**A)**

**0.87**

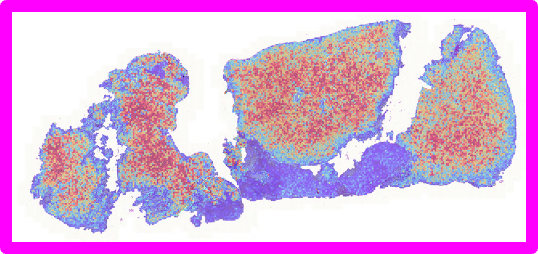

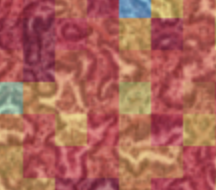

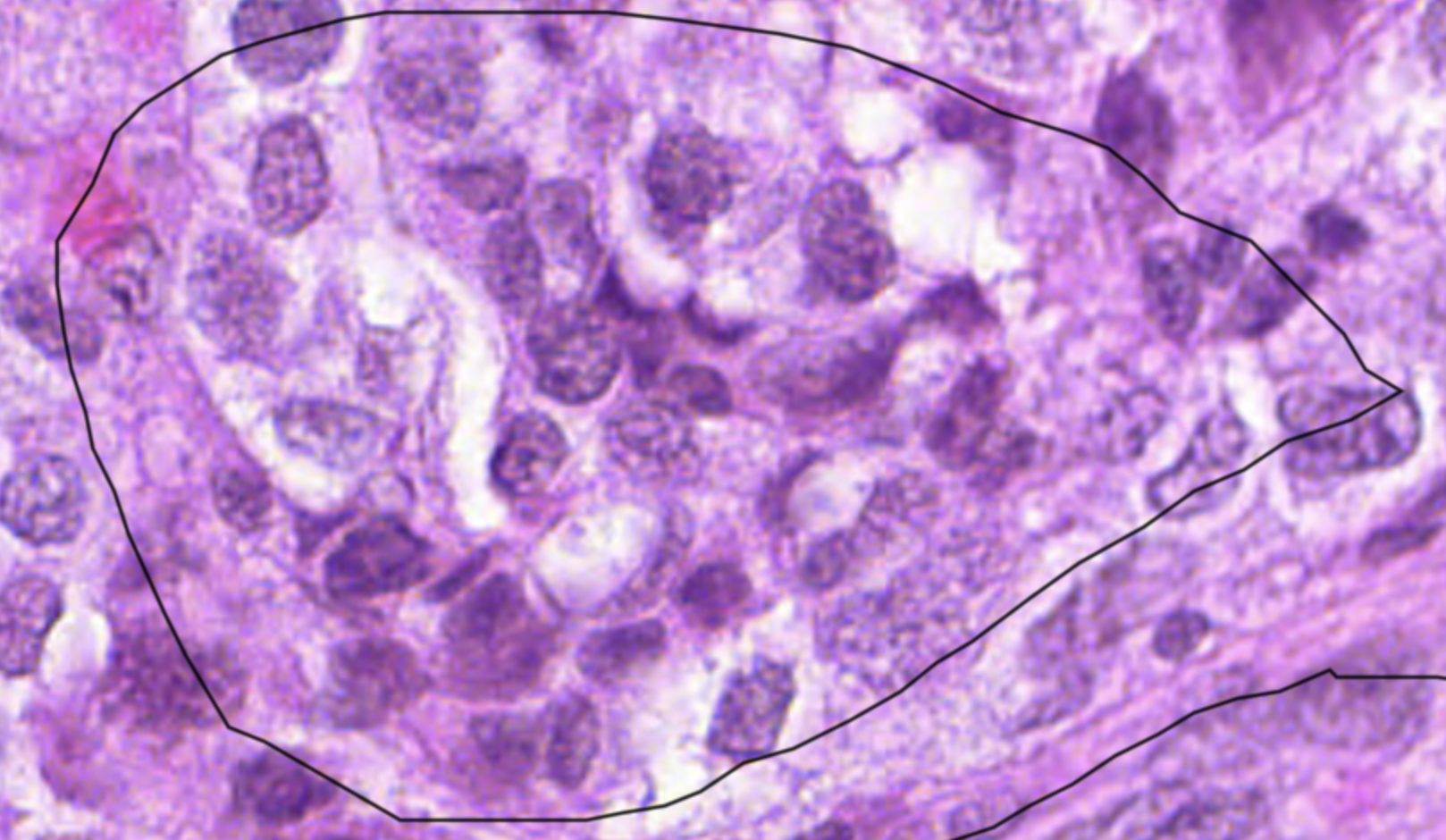

**B)**

**0.91**

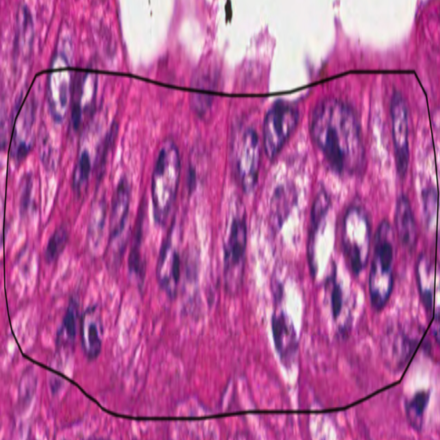

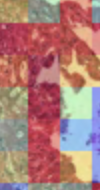

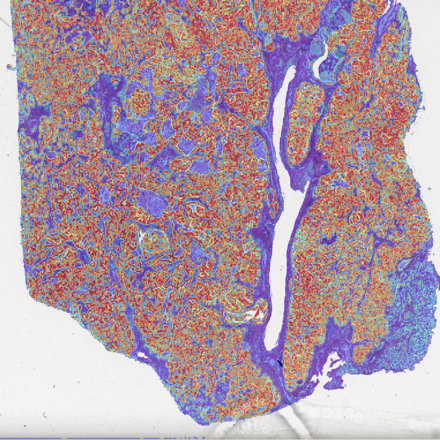

**Supplementary Figure S8**

**FOXO1**

A)

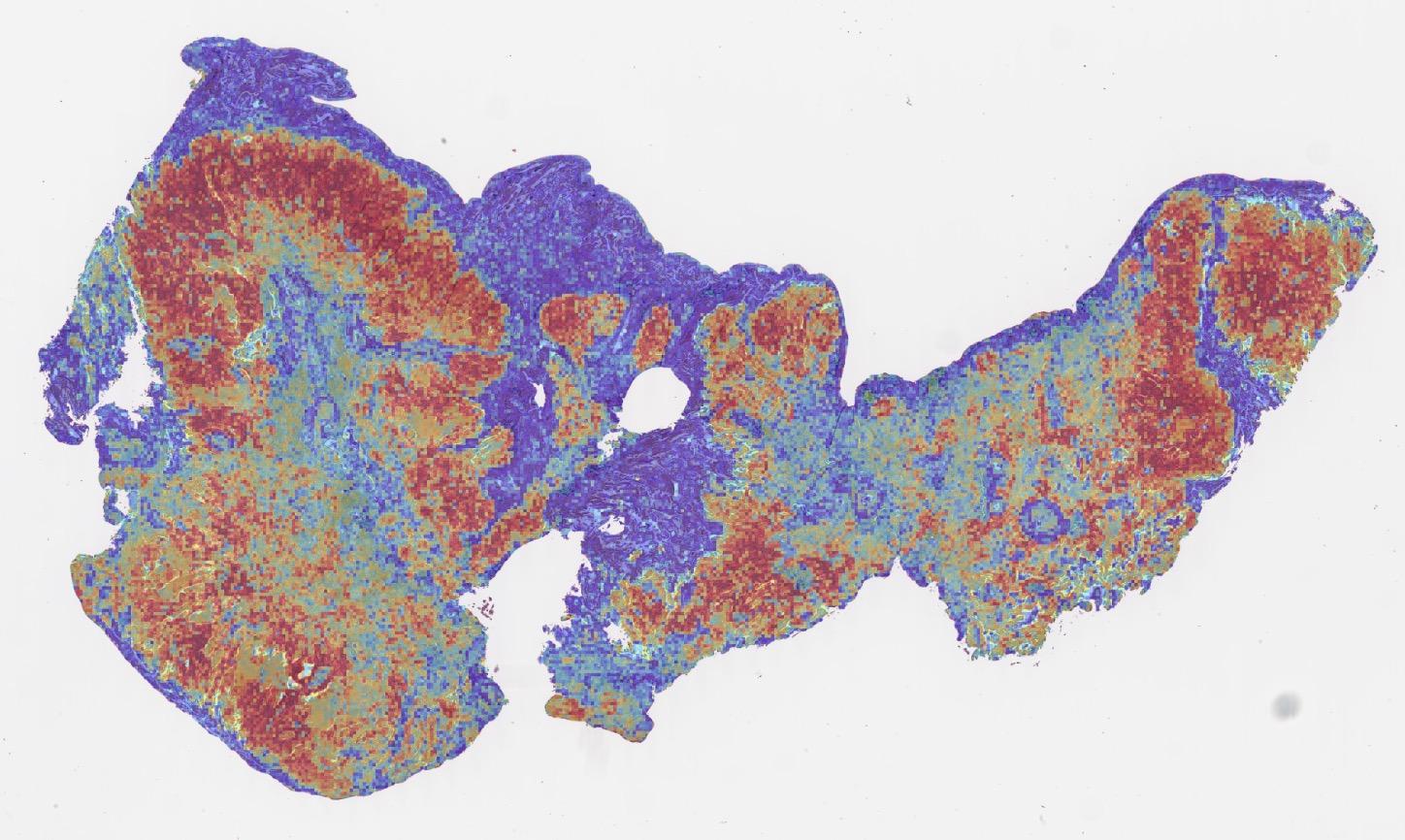

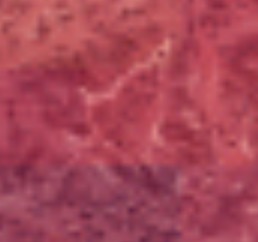

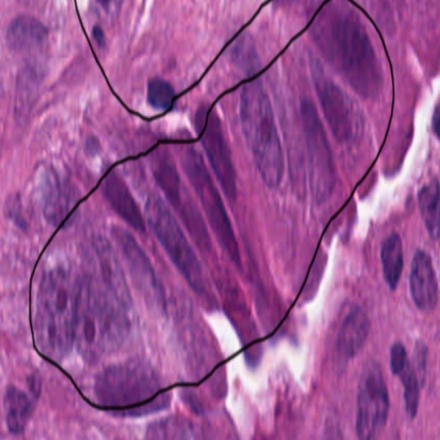

B)

**0.98**

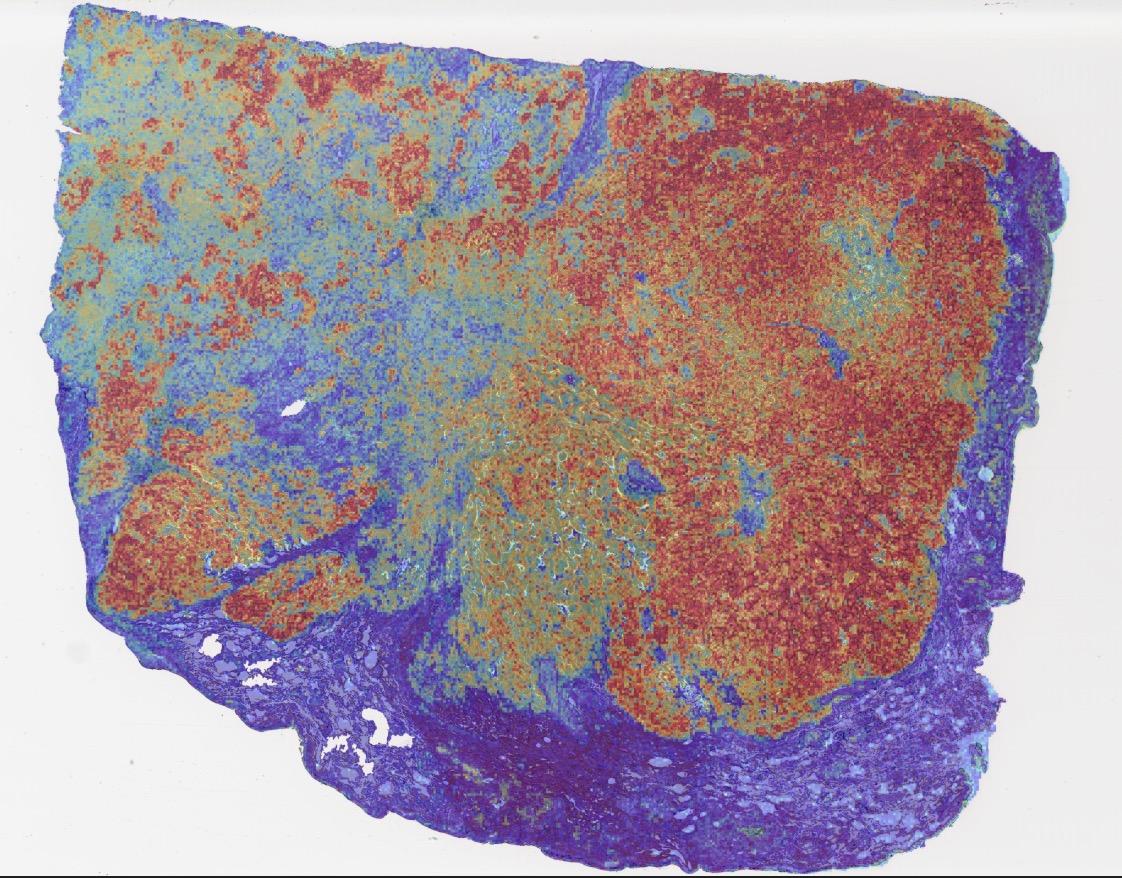

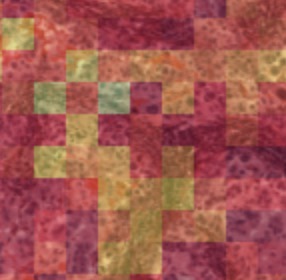

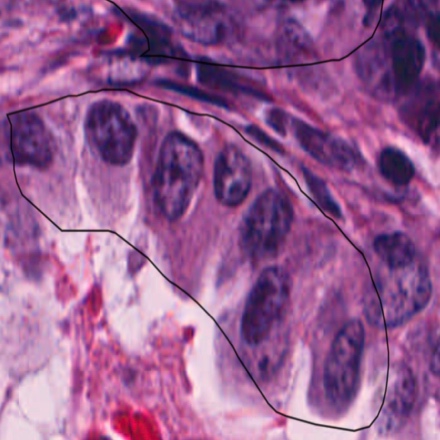

**0.85**

**Supplementary Figure S9**

**KEAP1**

A)

**0.96**

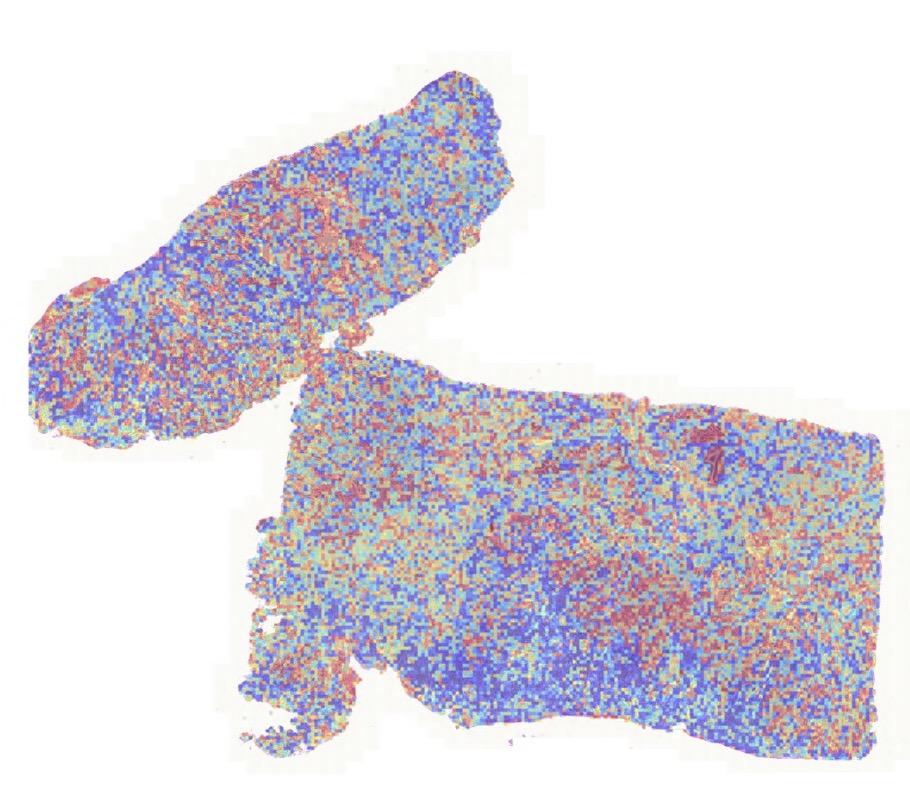

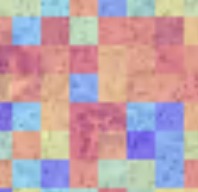

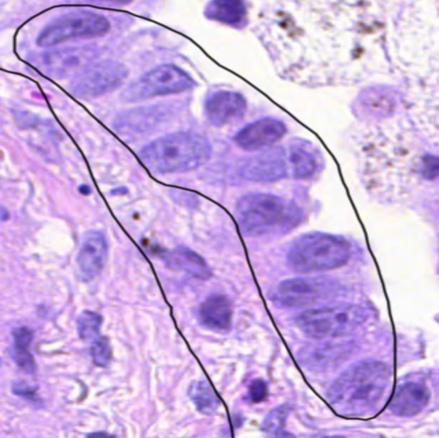

B)

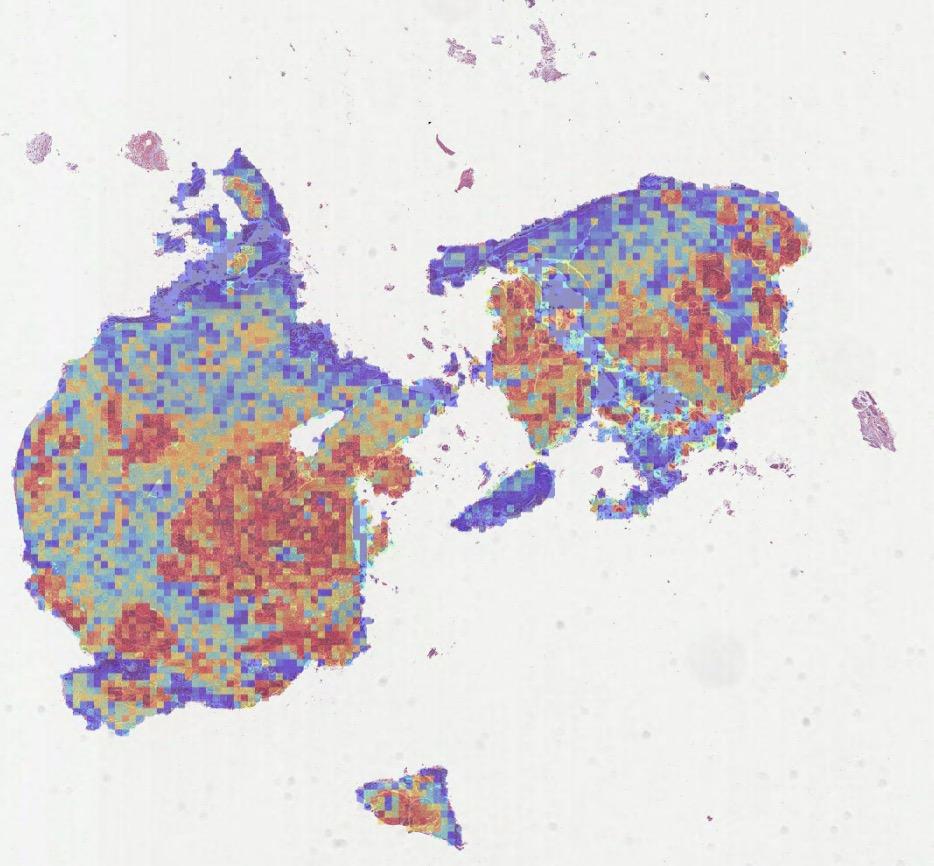

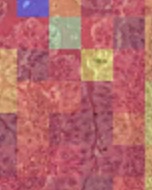

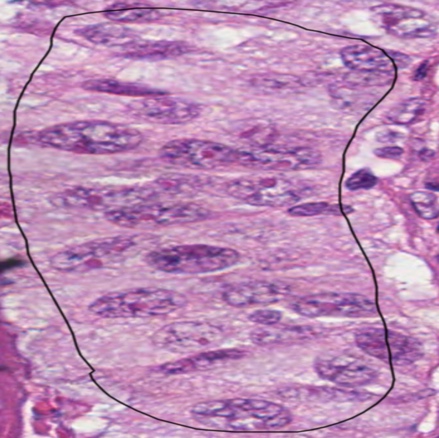

**0.98**

**Supplementary Figure S10**

**RB1**

A)

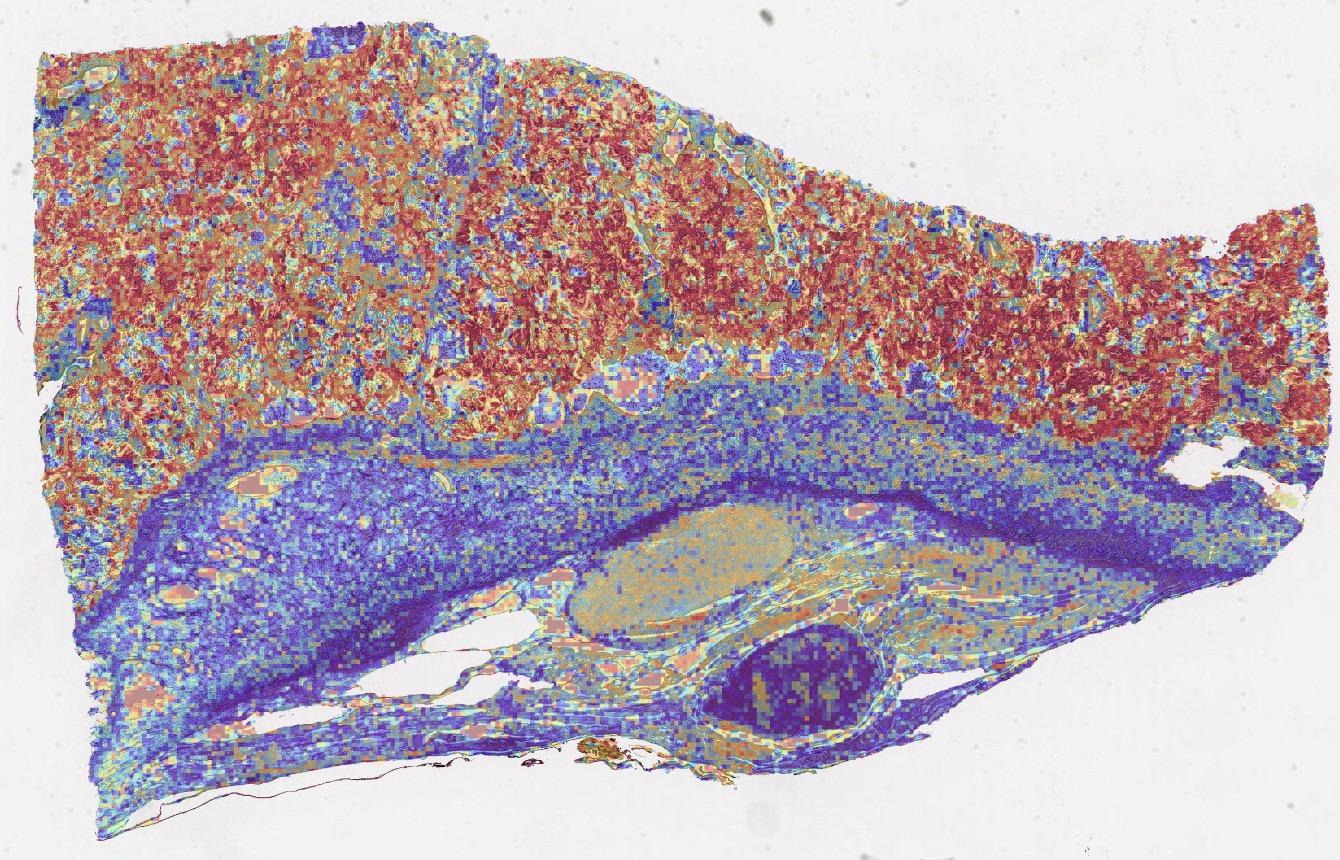

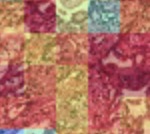

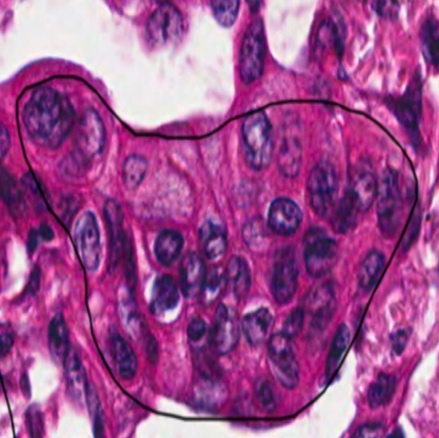

B)

**0.91**

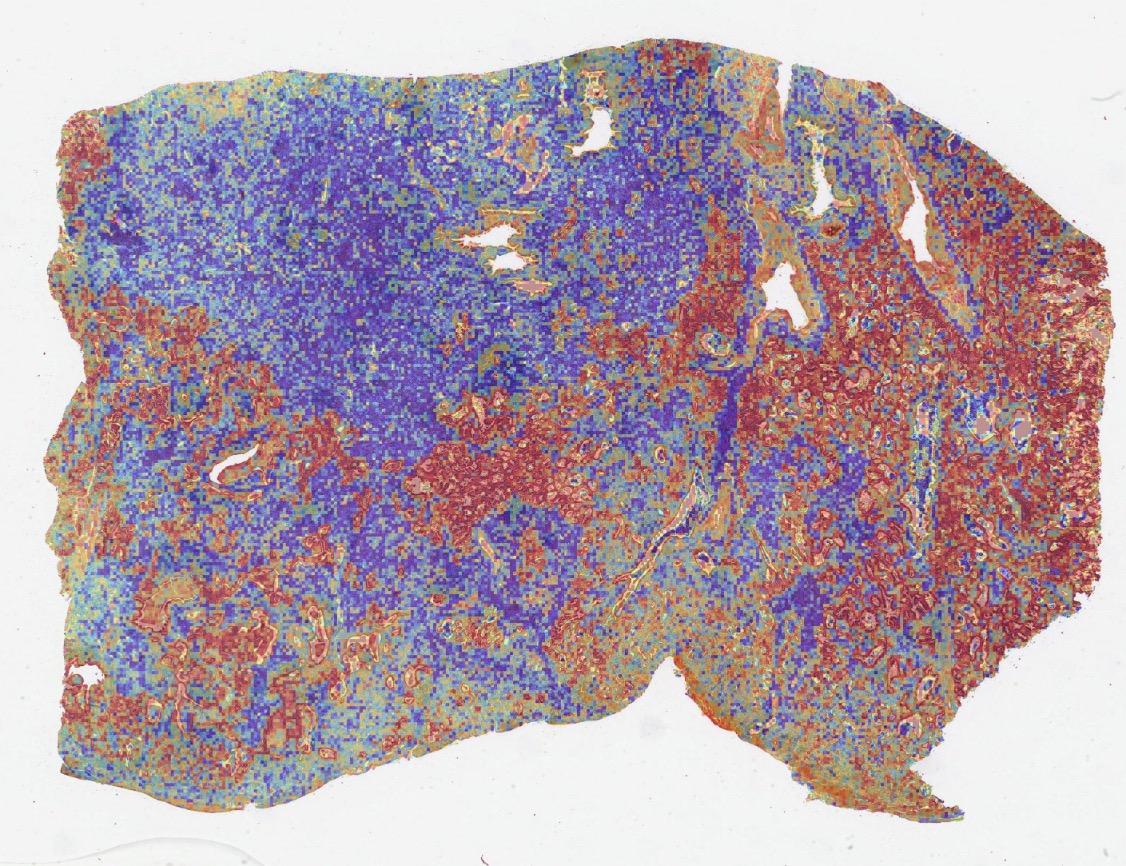

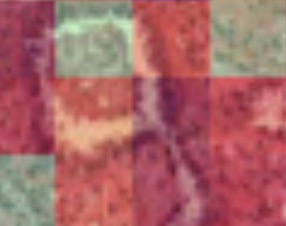

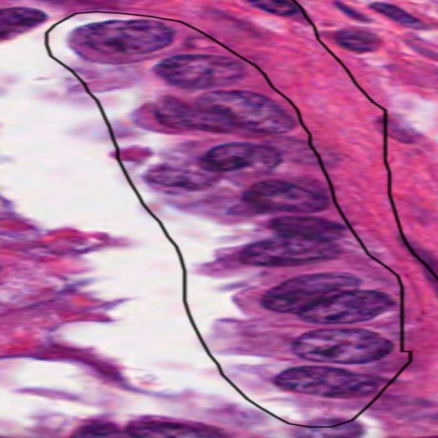

**0.92**

**Supplementary Figure S11**

**TP53**

A)

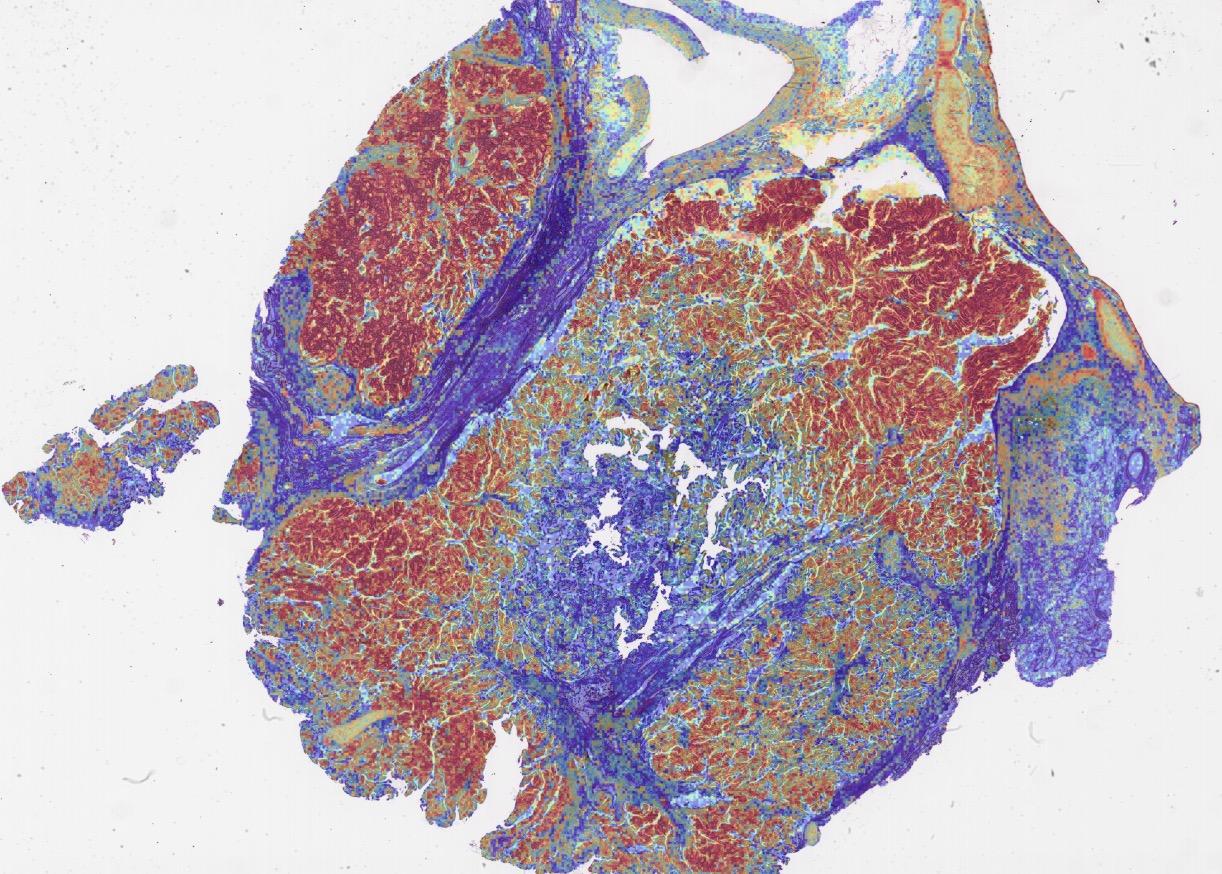

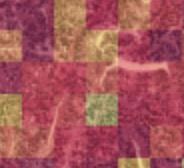

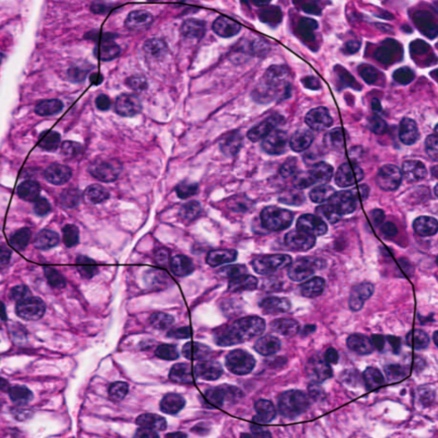

B)

**0.99**

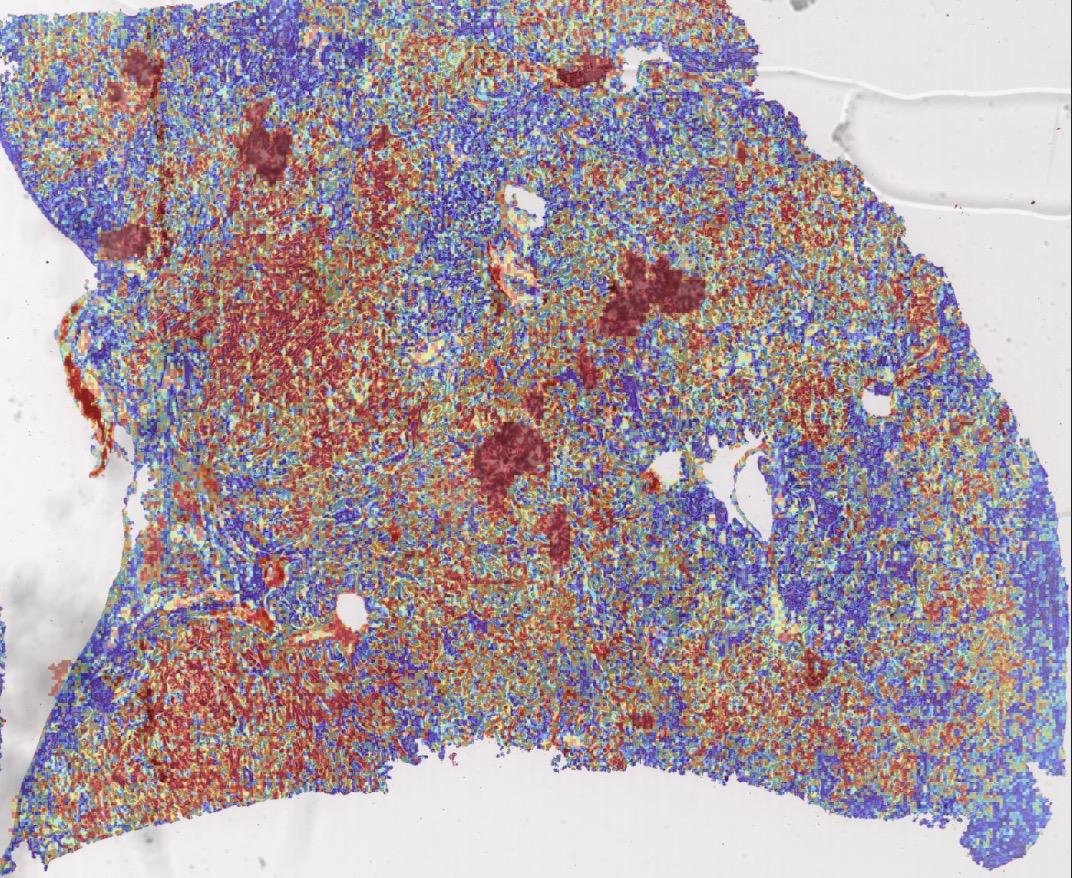

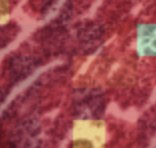

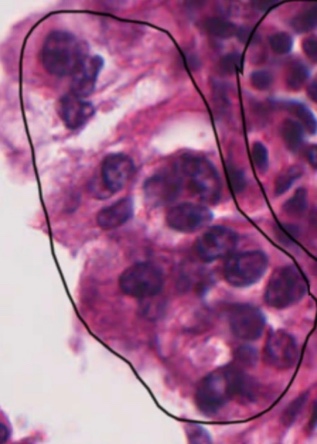

**0.97**

**Supplementary Figure S12**

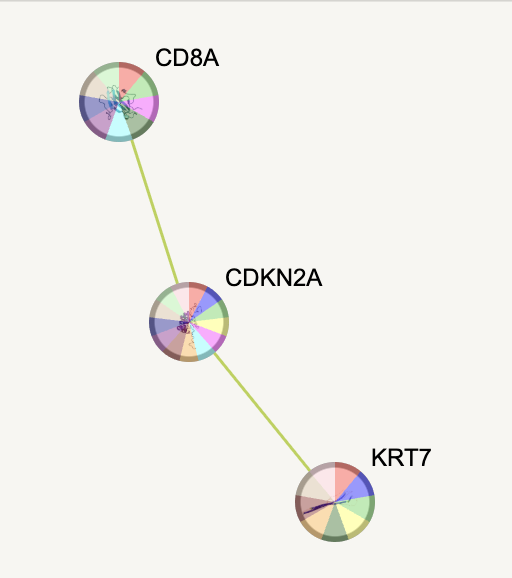

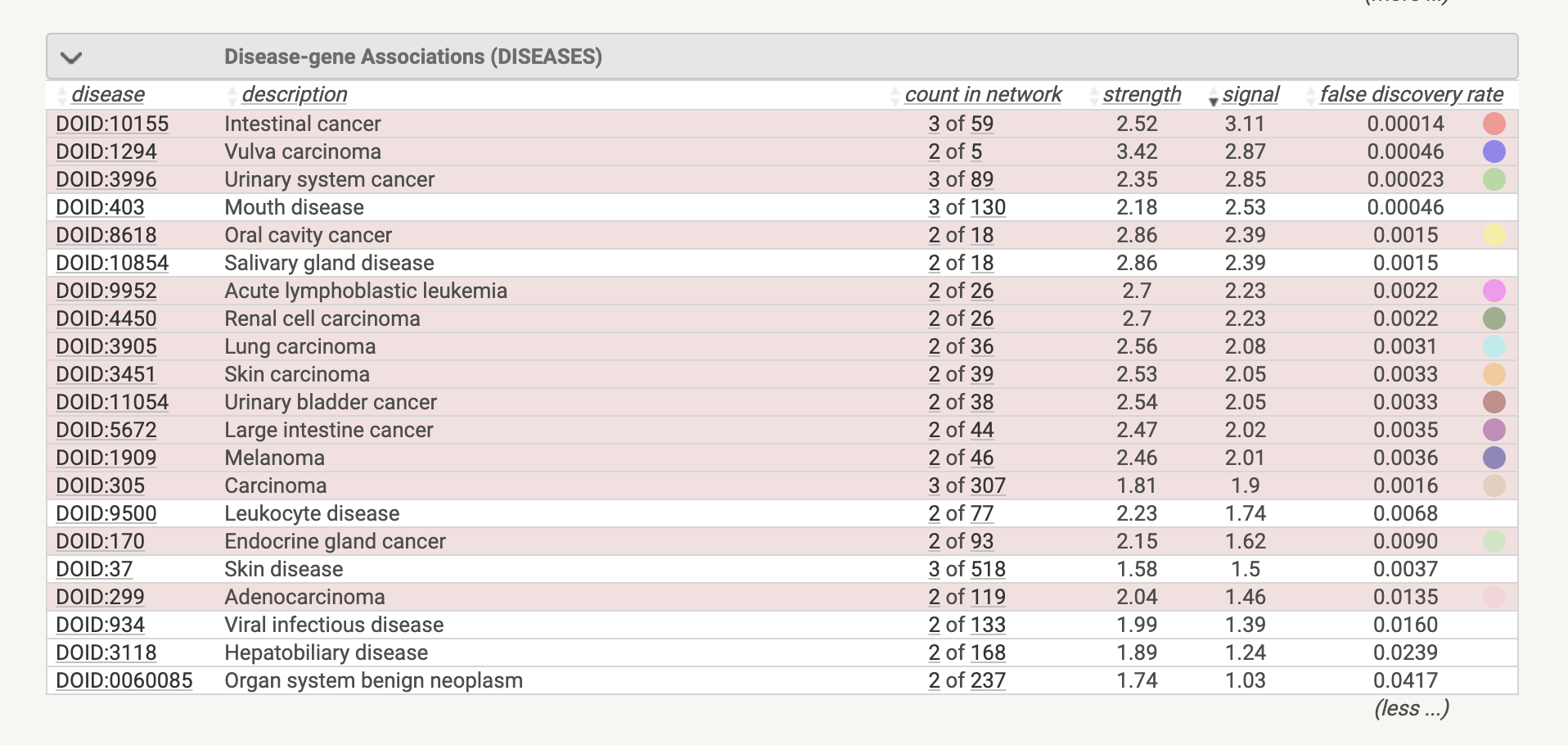

**Supplementary Figure S13-a**

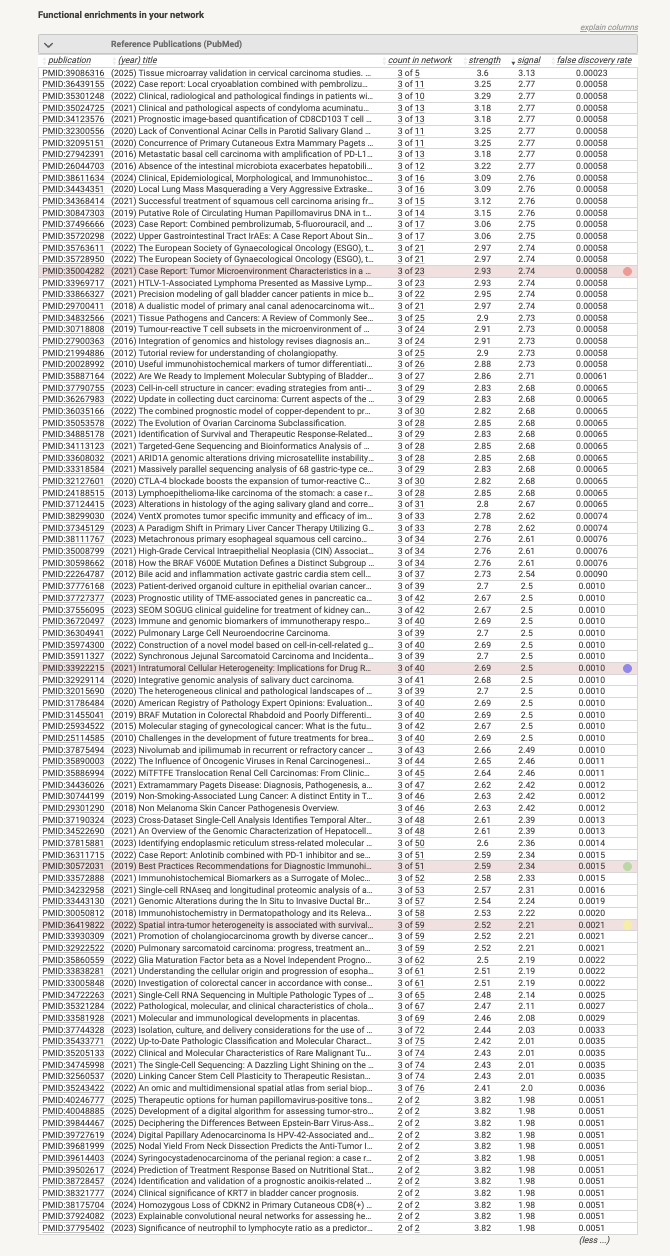

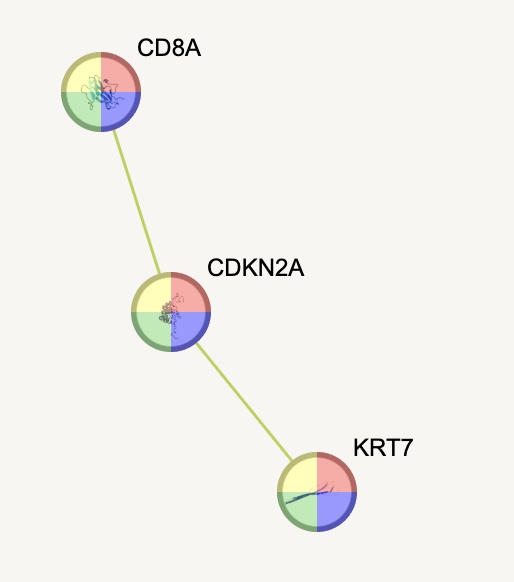

**Supplementary Figure S13-b**

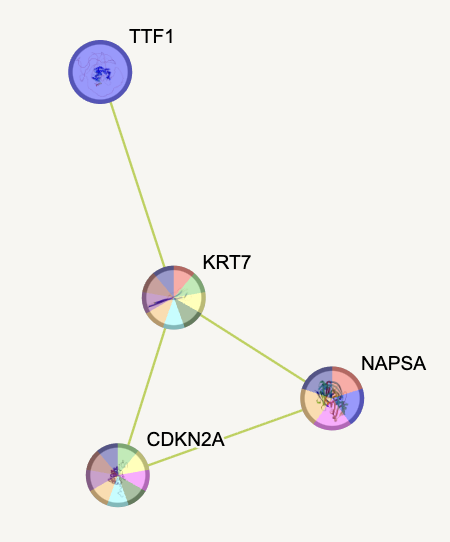

**Supplementary Figure S14-a**

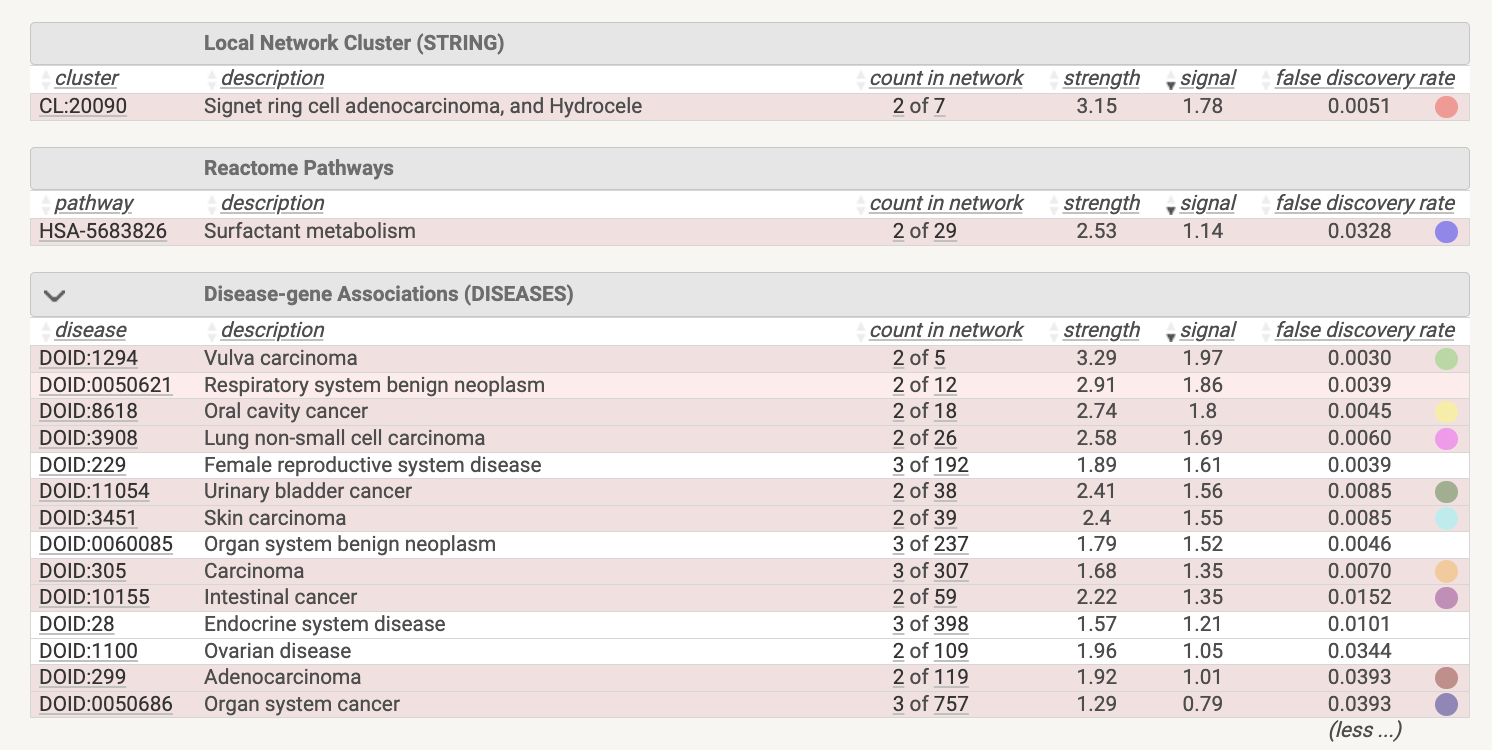

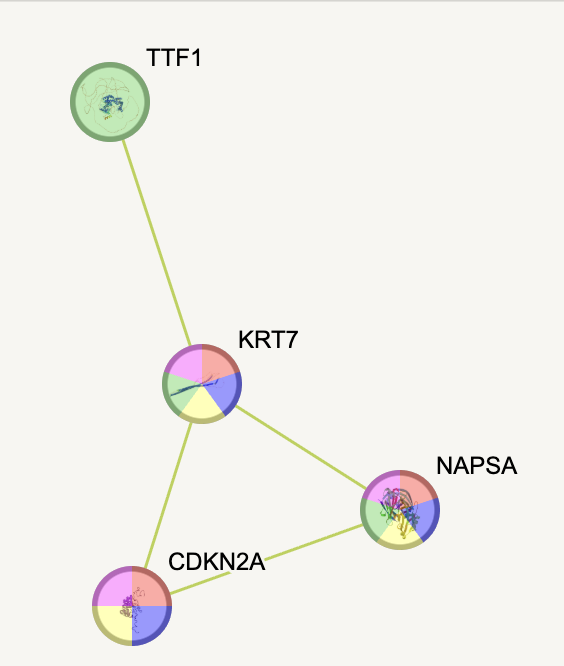

**Supplementary Figure S14-b**
